# Single-cell quantification of the microbiota by flow cytometry: MicFLY

**DOI:** 10.1101/2025.09.18.677130

**Authors:** Christine M. Tin, Bianca Cordazzo Vargas, Darryl A. Abbott, Aditi R. Lohar, Tiffany C. Taylor, Ivan A. Valishev, Sarah Weinshel, Vung Lian, Kacey J. Sullinger, Matthew Butoryak, Michael A. Silverman, Liat Shenhav, William H. DePas, Timothy W. Hand

## Abstract

The intestinal microbiota regulates multiple host functions, including digestion and immune development. Our knowledge of the microbiota has been shaped by available technology that primarily measures relative abundance. However, understanding the basis of shifts in microbiota composition requires single cell, absolute abundance measurements. In response to this problem, we developed <u>Mic</u>robiota <u>Fl</u>ow C<u>y</u>tometry (MicFLY), a single cell technology that directly quantifies and characterizes total bacterial abundances with species-level resolution in the microbiota. Using MicFLY, we can identify all major intestinal taxa, discriminate live from dead bacteria, perform single cell measurements of heterogeneous bacterial mRNA expression and concurrently quantify Immunoglobulin (Ig) A and G binding to intestinal bacteria. Using longitudinal species-resolved, quantitative analysis of the preterm infant microbiota, we identify that *E. coli* unbound by IgG and IgA associates with the development of necrotizing enterocolitis. The application of MicFLY single cell technology permits measurement of the microbiota at a finer scale and with deeper mechanistic understanding of compositional changes.

## INTRODUCTION

The human microbiome affects nearly all aspects of host physiology, and our technology has shaped how we understand these processes. A common method to measure the microbiome is 16S rRNA gene amplicon sequencing (16S-seq), which has allowed us to associate microbial diversity to a broad variety of host phenotypes and disease^1–5^. 16S-seq can be combined with the isolation of antibody bound-bacteria^6–8^ to measure relative immunogenicity of microbial members. In recent years, metagenomics has provided greater detail into the microbiome’s total gene content^9–11^, revealing functional differences between disease states^12^. However, these sequencing-based methods were developed for phylogenetic^13^ or functional^9^ classification, not the enumeration of different microbial species^14^. A limitation of sequencing is that its profiles are based on relative abundances, which is inadequate for describing population dynamics^15–17^. Absolute abundances are thus essential for interpreting the cause of compositional shifts. However, assays developed for this purpose^18–20^ ^21–24^ are subject to biases from PCR efficiency^25–30^ and DNA extraction^27,31–35^, potentially leading to distortions in microbial quantification. New technologies to quantify the microbiota with single cell resolution are necessary to identify the mechanisms by which host-microbiota interactions drive disease.

In this study, we present <u>Mic</u>robiota <u>Fl</u>ow C<u>y</u>tometry (MicFLY), a quantitative single cell technology that uses highly multiplexed bacterial probes and fluorescent hairpins to characterize and measure the microbiota. We verify that MicFLY can detect all major representative members of the microbiota at several taxonomic levels from phylum to species. Additionally, we show MicFLY can distinguish live from dead bacteria and measure taxon-targeted mRNA, thus linking transcript abundance to bacterial identity. By incorporating fluorescent antibodies, MicFLY can quantify host antibody binding to specific gut bacteria. Finally, we apply MicFLY to fecal samples collected from a longitudinal cohort of preterm infants and find that lack of Immunoglobulin G (IgG) and Immunoglobulin A (IgA) binding to *Escherichia coli* is associated with the subsequent development of necrotizing enterocolitis (NEC).

## RESULTS

### Microbiota Flow Cytometry (MicFLY): a single cell technique for measuring bacteria

MicFLY is a quantitative, single cell technology that uses high-dimensional spectral flow cytometry to measure individual bacteria in the microbiota (**Figure 1A**). Bacteria are first extracted from stool using centrifugation/density gradients to remove host cells and fecal debris. Isolated bacteria are stained with an amine-reactive fixable viability dye and anti-immunoglobulin fluorescent antibodies, which are then cross-linked to the cell surface in a two-step fixation protocol using bis(sulfosuccinimidyl)suberate (BS3) (a membrane-impermeable chemical crosslinker), followed by paraformaldehyde (PFA).

**Figure 1.**
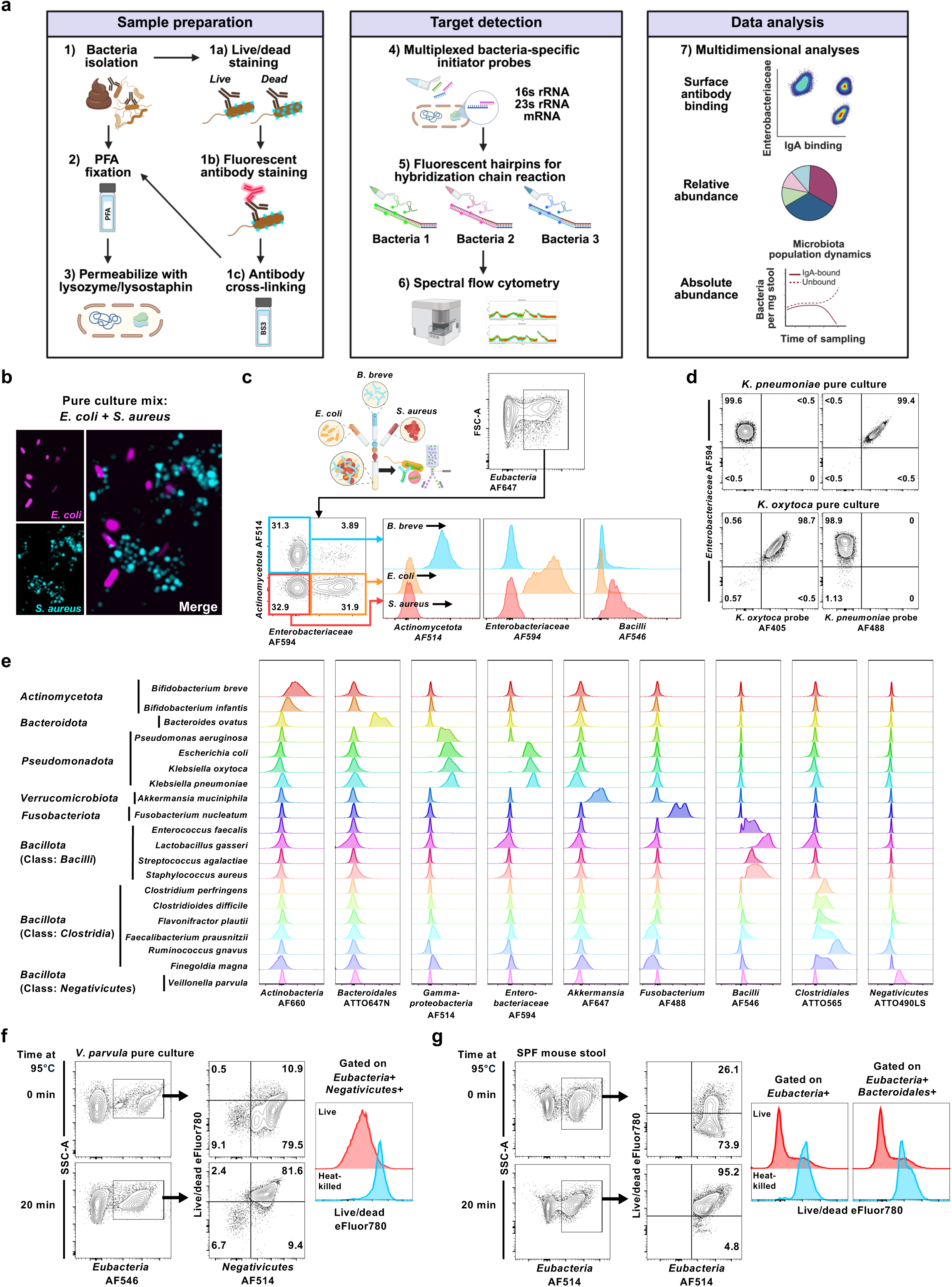
MicFLY single cell technology identifies diverse, live bacteria with high taxonomic resolution. **a,** Overview of MicFLY workflow (made with BioRender). **b**, Representative confocal microscopy images of *E. coli* (magenta; *Enterobacteriaceae* probe/B2-Alexa Fluor 594) and *S. aureus* (cyan; *Bacilli* probe/B4-Alexa Fluor 488) after staining by MicFLY protocol. **c**, Flow cytometry identification of *in vitro* grown bacteria in a mixed sample (*E. coli*, *S. aureus*, and *B. breve*) after MicFLY staining. Cells are gated on total bacteria (*Eub338* probe/B1-Alexa Fluor 647) prior to identification of individual taxa. Experimental design graphic made with BioRender. **d**, Discrimination of pure cultures of *K. pneumoniae* and *K. oxytoca* using species-specific probes by MicFLY. Cells are gated on total bacteria (*Eub338* probe/B1-Alexa Fluor 647). **e**, Multiplexed detection of 20 *in vitro* cultured bacterial species from diverse phyla representing the human gut microbiome by MicFLY staining. Each column represents signal from a unique probe across different taxa. All probes were tested in the same panel and stained on individual taxa. Cells are gated on total bacteria (as in **c**). **f**, Validation of viability dye on pure culture bacteria analyzed with MicFLY. *In vitro* cultured *Veillonella parvula* bacteria (*Negativicutes* probe/B5-Alexa Fluor 514) are heated to 95°C for 20 minutes prior to incubation with fixable viability dye eFluor780 and compared to non-heat killed sample. Flow cytometry plots (left) indicate gating strategy and histograms (right) indicate staining intensity with viability dye. **g**, Validation of viability dye on bacteria from fecal samples analyzed with MicFLY. Heat-killed and non-heat killed bacteria isolated from specific-pathogen free (SPF) C57BL/6 mouse stool are compared. Flow plots (left) indicate gating strategy and histograms (right) indicate viability dye staining to all bacteria (*Eubacteria* combined probe set/B5-AF514; center right) or *Bacteroidales* (*Bacteroidales* probe set/B17-Alexa Fluor 405; far right). All flow cytometry analyses show representative plots from multiple independent experiments. Numbers in plots represent percentage of events inside the gate. All histograms are normalized to mode.

Together, these fixatives increase the stability of fluorophore-conjugated antibodies bound to surface proteins and preserves intracellular architecture^36^. After fixation, bacteria are enzymatically permeabilized with lysozyme and lysostaphin, facilitating hybridization with oligonucleotide ‘initiator’ probes^37,38^ that target bacterial 16S and/or 23S ribosomal RNA (rRNA) and can be customized to taxa of interest **(Figure S1A, Table S1).** Probes targeting rRNA have an initiator sequence on their 3’ end that anneals to a unique pair of self-assembling fluorescent hairpins **(Figure S1B)**. Once annealed, hairpin pairs undergo a chain reaction (Hybridization Chain Reaction; HCR) to create a DNA scaffold capable of reaching over 10 kilobases^37–39^. Probe panel multiplexing is limited only by the number of hairpin-conjugated fluorophores available **(Supplemental Note S1, Table S2).** Importantly, by incorporating measurements of original sample mass with volumetric- or bead-based flow cytometry counting, absolute abundance can be calculated.

To generate taxon-specific initiator probes, we use both published sequences and those we designed with a curated rRNA database **(Table S1**, **Data S1).** In contrast to 16S-seq, MicFLY targets several sites along the entire rRNA molecule, enabling species-level resolution^40^. MicFLY also supports combinatorial logic with multi-fluorescent labeling of an individual bacterium at multiple taxonomic levels, massively improving identification capabilities.

A key advancement of MicFLY is the use of “*Eubacteria*” probes targeting conserved regions in the 16s rRNA bacterial subunit **(Figure S2A)**. As a proof-of-concept that *Eubacteria* probes discriminate bacteria from non-cellular debris, we tested the ‘universal’ probe (Eub338)^41^ on an arabinose-inducible Green Fluorescent Protein (GFP)-expressing *E. coli* strain^42^. Using GFP as a surrogate for *E. coli* presence, we demonstrated that virtually all GFP+ events were also stained with Eub338 **(Figure S2B)**. qPCR of GFP+ Eub338+ and GFP-Eub338-sorted fractions revealed *E. coli* ompX and 16S genes were undetectable in the Eub338-fraction, indicating false negatives are not a significant issue for MicFLY (**Figure S2B)**. Eub338-GFP-events also exhibit reduced light scatter properties **(Figure S2C)**, consistent with small particle debris^43,44^. As expected for two probes binding the same molecule^37^, comparison of Eub338 to a separate *Enterobacteriaceae* probe revealed a linear relationship **(Figure S2D).** While Eub338 is considered a universal bacterial marker, we designed two additional pan-bacterial initiator probes (Eub841 and Eub1491) to minimize the chance of undetected bacteria from the failure of any one probe **(Table S1, Data S2)**^45^.

Comparing these two novel probes reveal overlapping signal with Eub338 in mouse microbiota **(Figure S2E)**; critically, none of the probes bind to fecal matter from germ-free mouse stool, indicating lack of cross-reactivity with eukaryotic rRNA.

To test if MicFLY maintains cellular morphology, we stained and imaged mixed cultures of *Escherichia coli* (Gram-negative) and *Staphylococcus aureus* (Gram-positive) (**Figure 1B**). Confocal imaging revealed uniform morphology (*E. coli* = rods; *S. aureus* = cocci) and no ‘double’ stained bacteria, demonstrating that MicFLY is capable of measuring intact bacteria at the single cell level.

### MicFLY measures all major representative members of the intestinal microbiota at multiple taxonomic levels

We validated MicFLY can identify individual taxa in a mixed population using *in vitro* cultured *E. coli*, *S. aureus* and *Bifidobacterium breve*. Critically, MicFLY provided ‘digital’ identification of all three taxa, as each was positive only for their own probe (**Figure 1C**). To demonstrate MicFLY can differentiate between species, we designed probes targeting *Klebsiella pneumoniae* or *Klebsiella oxytoca*. Species-specific probes showed extraordinary specificity with virtually zero cross-reactivity (**Figure 1D**), demonstrating MicFLY can perform precise discrimination of closely related bacteria.

We designed MicFLY such that the fixation and permeabilization enable detection of bacteria with distinct cellular architecture. To test whether MicFLY was comprehensive to the most common bacteria in the human microbiota, we constructed a panel of ten initiator probes **(Table S1)** and assayed fixed cultures of 20 diverse species encompassing the dominant members of the human microbiota. MicFLY was able to identify each taxon with high specificity and minimal off-target signal (**Figure 1E**), highlighting that MicFLY is effective on a broad variety of bacteria.

### MicFLY quantifies the number of live bacteria

Flow cytometry has been used to measure total bacterial abundance^46–50^, typically using nucleic acid stains that are not exclusive to bacteria and lack taxon specificity. To establish MicFLY’s capacity for bacterial quantification, we serially diluted four bacteria (*E. coli*, *K. pneumoniae*, *E. faecalis*, and *S. aureus*) and compared abundances in parallel across three methods: 1) MicFLY; 2) DNA staining (SytoBC), and 3) colony formation **(Figure S3A).** Using colony-forming units (CFU) as the ‘gold standard’ comparison, MicFLY quantification was superior to nucleic acid staining, showing a linear relationship over a wider range (∼10^4^-10^8^ counts/mL for MicFLY vs. ∼10^6^-10^8^ counts/mL for SytoBC) and closer adherence to CFU for all four taxa, as calculated by cumulative perpendicular distance **(Figure S3A).** We also evaluated MicFLY’s ability to enumerate bacteria from preterm stool **(Figure S3B)**. Though CFU counts cannot be performed for fecal samples, we found MicFLY was comparable to Syto BC between ∼10^4^-10^7^ counts/mL **(Figure S3C)**. No correlation was observed between MicFLY-calculated total abundance and input stool, revealing mass is a poor surrogate in such samples for bacterial content **(Figure S3D)**.

Current microbiota analysis methods are inefficient for distinguishing viable bacteria^51,52^, hindering our ability to study the important but distinct roles that live and dead bacteria play in host physiology^53–59^. To address this, we tested an amine-reactive fixable viability dye in our MicFLY protocol and showed clear discrimination of viable bacteria in heat-killed bacterial culture (**Figure 1F**) and mouse stool (**Figure 1G**). Overall, MicFLY functions well for single cell quantification of the live microbiota.

### Validation of MicFLY *in vivo* with a known bacterial community

We next validated MicFLY for microbiota identification and quantification using gnotobiotic mice colonized with a predefined 9-member community of bacteria derived from young mice (Pediatric Community; PedsCom)^60,61^ (**Figure 2A**) and compared relative/absolute abundances between MicFLY, qPCR and 16S-seq. To have comparable taxonomic resolution to 16S-seq, we generated family-level probes (7 different families) and validated each on pure culture isolates of each bacterial member **(Figure S4A)**. Relative abundance measurements showed high concordance between MicFLY and 16S-seq (**Figures 2B** and **2C**). Accordingly, Bray-Curtis beta-diversity distance analysis confirmed that relative composition measured by MicFLY and 16S-seq was more similar to each other than to qPCR (**Figure 2C**). Absolute abundance counts demonstrated that total numbers for the majority of bacteria were comparable between MicFLY and qPCR, except for the *Lactobacillaceae* which MicFLY counted as significantly higher (**Figure 2D**). Enhanced detection of this family was also reproduced in a smaller gnotobiotic community (PedsCom-6)^61^ (**Figure S4B**). Bacterial abundances calculated by MicFLY were not artificially increased by background auto-fluorescent events, as germ-free mouse stool established that these events (∼10^1^-10^3^ cells/mg stool) are too rare to have an impact on abundance measurements **(Figure S4C).**

**Figure 2.**
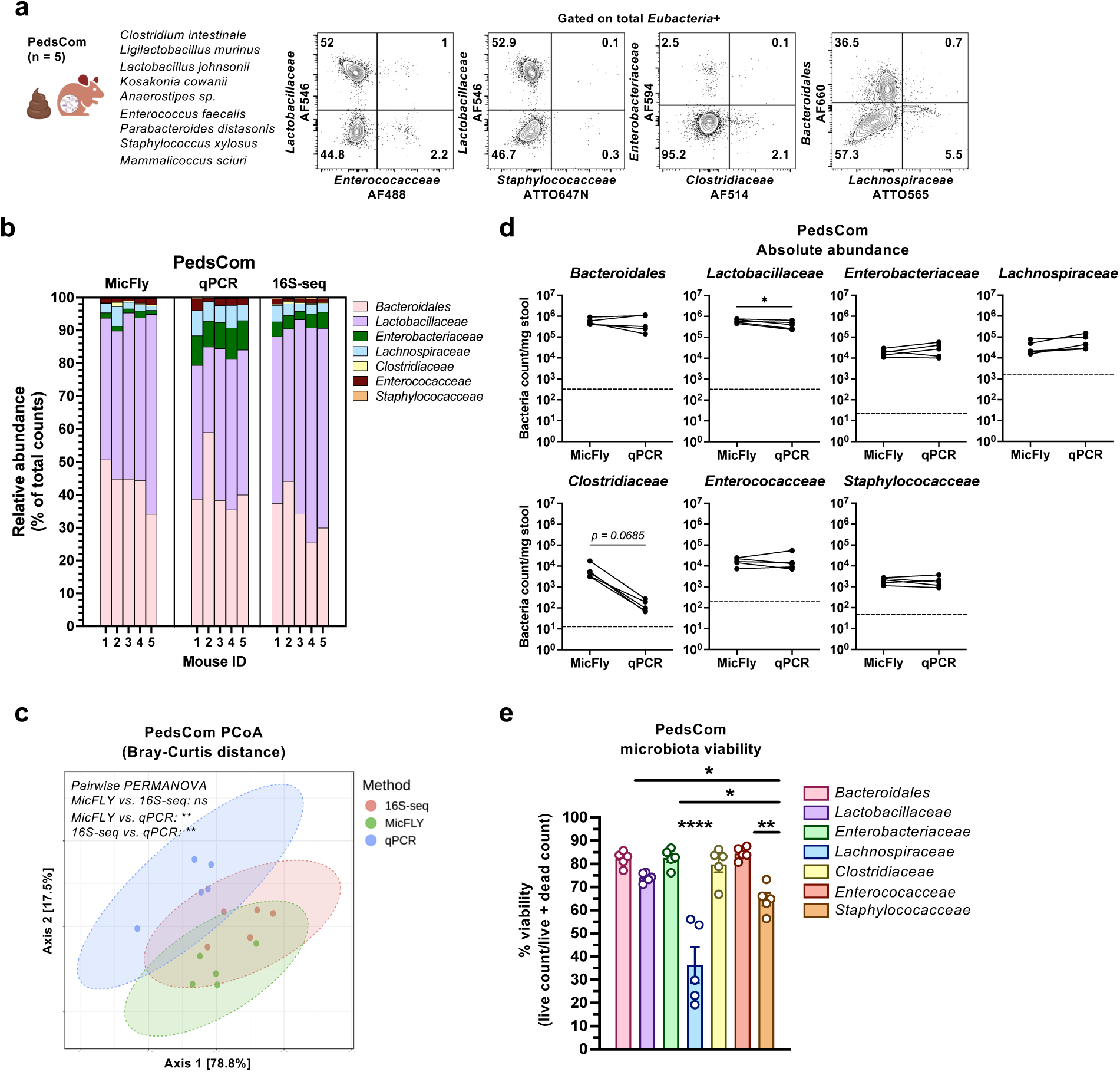
MicFLY quantification and viability assessment of a predefined microbial consortia. **a**, MicFLY staining and spectral flow cytometry resolves all taxa within a defined bacterial community. Fecal samples were isolated from gnotobiotic Eα16/NOD mice (n = 5; ages 4-20 weeks old) colonized with a nine-member (PedsCom) bacterial community (described in image; left). Shown are representative flow plots from MicFLY analysis at the family-level, gated on total bacterial cells using the *Eubacteria* combined probe set/B17-Alexa Fluor 405. **b**, The composition of PedsCom fecal samples was analyzed by MicFLY, quantitative PCR and 16S rRNA gene amplicon sequencing (16S-seq; V4 region). Stacked bar charts depict relative abundance calculated by each method. To calculate relative abundance for qPCR, the number of counts for a taxon was divided by the sum of counts for all measured taxa. PedsCom relative abundance measured by MicFLY was calculated the same way. 16S-seq relative abundance data were processed using QIIME2. **c**, Principal coordinates analysis (PCoA; Bray-Curtis using MicrobiomeAnalyst^127^) depicts beta diversity measurements for each microbiota measurement method. Ellipses represent 95% confidence intervals for each group, and statistical comparisons made using pairwise PERMANOVA with multi-testing adjustments based on Benjamini-Hochberg (FDR). p < 0.001 (**); ns, not significant (p ≥ 0.05) **d**, Absolute abundances of each bacterial family measured by MicFLY and qPCR from paired stool samples for five mice. Bacterial counts normalized to stool weight (mg), and statistical comparisons performed using a paired two-tailed t-test. Dashed line represents limit of detection for that taxon, calculated as mean absolute abundance in stool from germ-free mice (n = 4). p < 0.05 (*). **e**, Microbiota viability as measured by MicFLY. Statistical significance assessed using one-way ANOVA with Tukey’s multiple comparisons test. Bars show mean± SEM. p < 0.05 (*), p < 0.01 (**), p < 0.0001 (****). Group with four asterisks is significantly different from every other group. DNA extracted from one stool pellet for each mouse (five mice total) was used for both qPCR and 16S-seq. A second stool pellet from each mouse was tested by MicFLY. For MicFLY and qPCR, each pellet was tested twice in two independent runs, and data reflect the average relative/absolute abundance **(b-d)** or average viability **(e)** of both runs. For 16S-seq, data reflect relative abundances from a single sequencing experiment **(b, c).**

For these comparative abundance analyses, we measured total *Eubacteria*+ cells by MicFLY (both live and dead) to maintain consistency with bulk DNA tests (**Figures 2A-2D)**. We were also interested in whether different taxa were over- or under-represented by live or dead cells. We quantified the live microbiota **(Figure S4D)** and uncovered that the majority of taxa had similar viability (∼80%), except for *Lachnospiraceae* and *Staphylococacceae* which were significantly less viable (**Figure 2E**). Re-analysis of absolute abundance data focusing on live bacteria revealed that qPCR overestimated the number of viable *Lachnospiraceae* **(Figure S4E)**. These findings uncover how bulk cell analyses may be substantially affected by dead bacteria in some microbial communities, which can be circumvented using MicFLY which provides single cell live/dead discrimination.

### MicFly measures bacterial mRNA with single cell resolution

MicFLY uses HCR, which is effective for detecting bacterial gene expression^37,62–69^, but has not been used to measure taxon-targeted mRNA in the microbiota. To measure mRNA with MicFLY, we used a split-initiator design in which two initiator probes must anneal to their target at adjacent sites for fluorescent hairpins to unfold **(Figure S1B)**^37,70^. We targeted multiple sites along the mRNA to maximize fluorescent signal **(Table S3).** To assess whether MicFLY preparation conditions maintain detectable levels of mRNA, we tested *E. coli* transformed with a plasmid containing GFP controlled by a pBAD arabinose-inducible promoter^42,71,72^ (**Figure 3A**). Immediately post-induction, MicFLY measured a rapid rise in both GFP mRNA and protein **(Figures S5A and S5B).** After four hours, GFP mRNA dropped while reporter protein remained high. This reduction in mRNA was not due to arabinose depletion as ‘spent media’ could induce GFP from unstimulated *E. coli* **(Figure S5C)**. These results are consistent with negative feedback occurring in the pBAD system due to self-regulation from arabinose metabolism products^73–75^.

**Figure 3.**
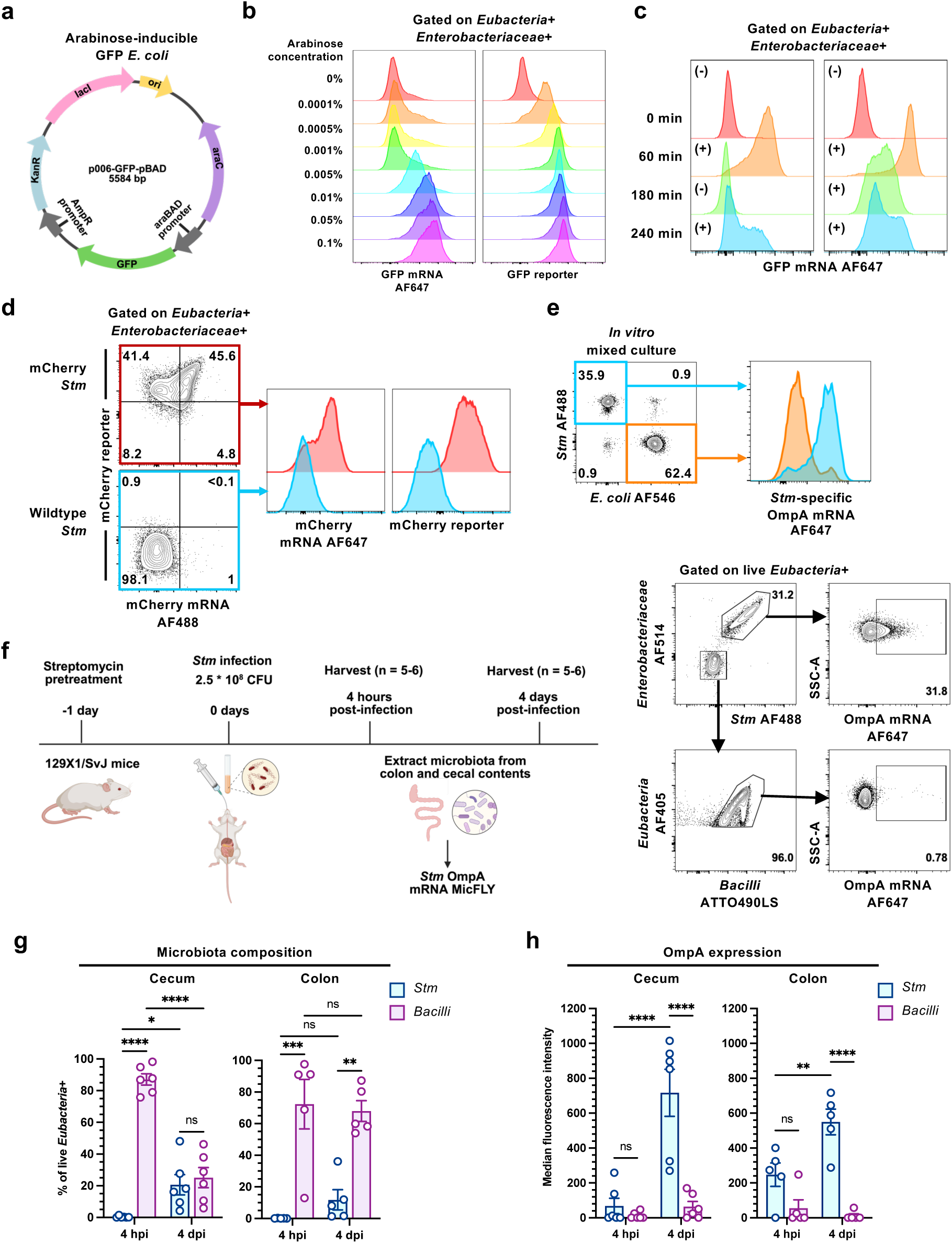
MicFLY enables single-cell quantification of bacterial mRNA expression *in vitro* and *in vivo*. **a**, Plasmid map of pBAD promoter-driven arabinose-inducible GFP mRNA with kanamycin resistant gene. **b**, GFP mRNA and protein levels in *E. coli* induced with increasing concentrations of arabinose (% w/v), measured with MicFLY. mRNA-targeted probes were used to detect GFP transcripts, while GFP reporter protein was directly measurable by flow cytometry. **c**, Dynamics of GFP mRNA expression over time in *E. coli* following arabinose induction (+) or removal (−), measured with MicFLY. **d**, Flow cytometric detection of mCherry mRNA and mCherry reporter protein in a *Salmonella enterica serovar Typhimurium* (*Stm*) strain carrying a genomically-encoded, constitutively expressed mCherry reporter protein, measured with MicFLY. *Stm* strain lacking mCherry was used as a control. mCherry transcripts were detected by MicFLY mRNA-targeted probes, while mCherry reporter protein was directly measurable by flow cytometry. **e**, Detection of *Stm* ompA mRNA in a mixed sample of *in vitro* cultured *Stm* and *E. coli.* **f**, Experimental design (left; made with BioRender) of *in vivo* ompA mRNA expression detection. 129X1/SvJ adult mice (n = 5-6 per time-point) were treated with streptomycin, then orally infected the next day with 2.5 × 10 CFU *Stm*. *Stm*-infected mice were sacrificed at either 4 hours post-infection (4 hpi) or 4 days post-infection (4 dpi) for MicFLY analysis of cecal and colonic contents. Representative flow plots (right) depict gating scheme and ompA mRNA expression for *Stm* or *Bacilli*. **g**, Microbiota composition in cecum and colon at 4 hpi and 4 dpi as measured by MicFLY. Graphs depict relative proportions of live *Eubacteria+Stm+* and live *Eubacteria*+*Bacilli+* populations. Bars show mean ± SEM. Each dot represents an individual mouse. **h**, Graphs depict the median fluorescence intensity (MFI) of ompA mRNA expression in live *Eubacteria+Stm+* or live *Eubacteria*+*Bacilli+* populations at 4 hpi and 4 dpi. Bars show average MFI ± SEM. Each dot represents an individual mouse. For *in vitro* tests **(b-e)**, bacteria were gated on *Eubacteria*/B17-Alexa Fluor405+ *Enterobacteriaceae*/B2-Alexa Fluor 594+. Numbers in flow plots represent percentage of events inside the gate, and all histograms are normalized to mode. For *in vivo* experiments **(f-h),** two independent sets of mice were tested (set 1: *n=3*; set 2: *n=2-3*). Set 1 used probes for *Eubacteria, Enterobacteriaceae, Stm, Lactobacillaceae* (family-level), and ompA. Set 2 used probes for *Eubacteria, Enterobacteriaceae, Stm, Bacilli* (class-level), and ompA. For statistical analyses, *Lactobacillaceae* and *Bacilli* data were combined and reported as *Bacilli*. Statistical tests were performed using two-way ANOVA with Tukey’s multiple comparisons test. p < 0.05 (*), p < 0.01 (**), p < 0.001 (***), p < 0.0001 (****).

To assess MicFLY’s sensitivity to different mRNA levels, we tested varying arabinose concentrations for induction (**Figures 3B and S5D)**. Messenger RNA levels were more sensitive to arabinose concentration than GFP protein, likely because of the short half-life of prokaryotic mRNA^76^. To track kinetic differences in mRNA expression, we added and removed arabinose at different time points and compared transcript level to that with a constant arabinose supply. MicFLY could identify heterogeneous subpopulations that emerged at different intervals **(Figure S5E)** and measure a dynamic transcriptional response (**Figure 3C**). These findings are consistent with heterogeneous on-switching of the pBAD promoter, which results in both induced and non-induced populations at any given time^73,77,78^, captured here at the single cell level.

Messenger RNA levels produced by high-copy plasmids can be artificially elevated. Thus, it was important to test whether MicFLY could also measure mRNA expressed from the chromosome. To test this idea, we measured a constitutively-expressed, genomically-encoded mCherry mRNA and protein in a strain of *Salmonella enterica serovar Typhimurium* (*Stm*)^79^ (**Figure 3D**). Here we observed that mCherry protein was consistently high while mRNA was expressed bimodally, indicating heterogeneity in mRNA levels. Collectively, these analyses demonstrate MicFLY resolves mRNA transcript levels in single cells *in vitro* and thus can identify heterogeneous subpopulations with asynchronous gene expression.

Being able to concurrently identify and measure mRNA from individual taxa in mixed populations would be a major advancement. To determine whether MicFLY could perform this function, we measured mRNA expression of the *Stm* outer membrane protein (ompA) and confirmed its detection in *Stm* in a mixed population with *E. coli* (**Figure 3E**). As the ompA gene is present in both *Stm* and *E. coli* and only differs by ∼10% nucleotide identity **(Figure S5F)**, this result demonstrates the accuracy and precision of MicFLY in discriminating specific mRNAs. We next measured ompA expression *in vivo* using a common intestinal infection model of *Stm*^80,81^ (**Figure 3F**). We identified *Stm* and *Bacilli* as major constituents in the cecum and colon (**Figure 3G**), and found that ompA expression levels in *Stm* increase significantly at later time points of infection, but is not detected in *Bacilli* (**Figure 3H**). These data demonstrate that MicFLY can perform quantitative, targeted, single-cell transcriptomics in the gut microbiota.

### MicFLY quantifies taxonomically resolved IgA-bound single bacterial cells

We designed the MicFLY fixation and permeabilization protocol to maintain the structure of proteins, such as host antibodies, on the bacterial surface. To test this, we incubated *E. coli* with breast milk IgA according to our published protocol^82^. Comparison of IgA-bound fixed bacteria before and after MicFLY showed no difference in binding (**Figure S6A**). To assess the specificity of antibody binding in MicFLY, we analyzed bacteria extracted from the stool of wildtype (C57BL/6) or IgA^-/-^ mice and observed virtually no IgA staining in IgA^-/-^ samples (**Figures S6B and S6C**). In wildtype mice, IgA binding to various taxa was not uniform, as we found that the *Lactobacillaceae* were highly IgA-bound compared to other bacteria **(Figure S6C**). Thus, MicFLY now makes possible the simultaneous identification and quantification of IgA-bound taxa at the single cell level from mixed populations collected *in vivo*.

### MicFLY identifies altered IgA binding patterns in donor milk-fed preterm infants

Preterm infants are colonized with an atypical microbiota dominated by facultative anaerobic bacteria (*Enterobacteriaceae*, *Bacilli*), which is associated with increased incidence of diseases like NEC^83^. Development of the infant microbiota (including preterm) is regulated by feeding modality with bioactive molecules in milk (IgA, oligosaccharides etc.) as potent mediators^84^. Human milk is associated with reduced incidence of NEC^85^, and when the mother’s own milk (MOM) is not available, donated human milk is often used as a substitute. Donor milk (DM) must be pasteurized to reduce the risk of infection, but pasteurization reduces the amount^86–88^ and anti-bacterial reactivity of IgA^82^. How much milk-derived IgA is required to saturate the infant microbiota, and whether DM leads to reduced IgA binding of intestinal bacteria, remains unknown. To test this, we used MicFLY to analyze fecal samples from a longitudinal prospective cohort study of preterm infants born between 24 and 33 gestational weeks of age, with stool collection starting at day of life (DOL) 10 (**Figure 4A**). We selected samples from infants that i) were not treated with antibiotics after DOL 7, ii) did not develop NEC or other infection-related illnesses, and iii) were either fed exclusively MOM (n = 6 infants, 29 samples) or exclusively DM (n = 12 infants, 44 samples).

**Figure 4.**
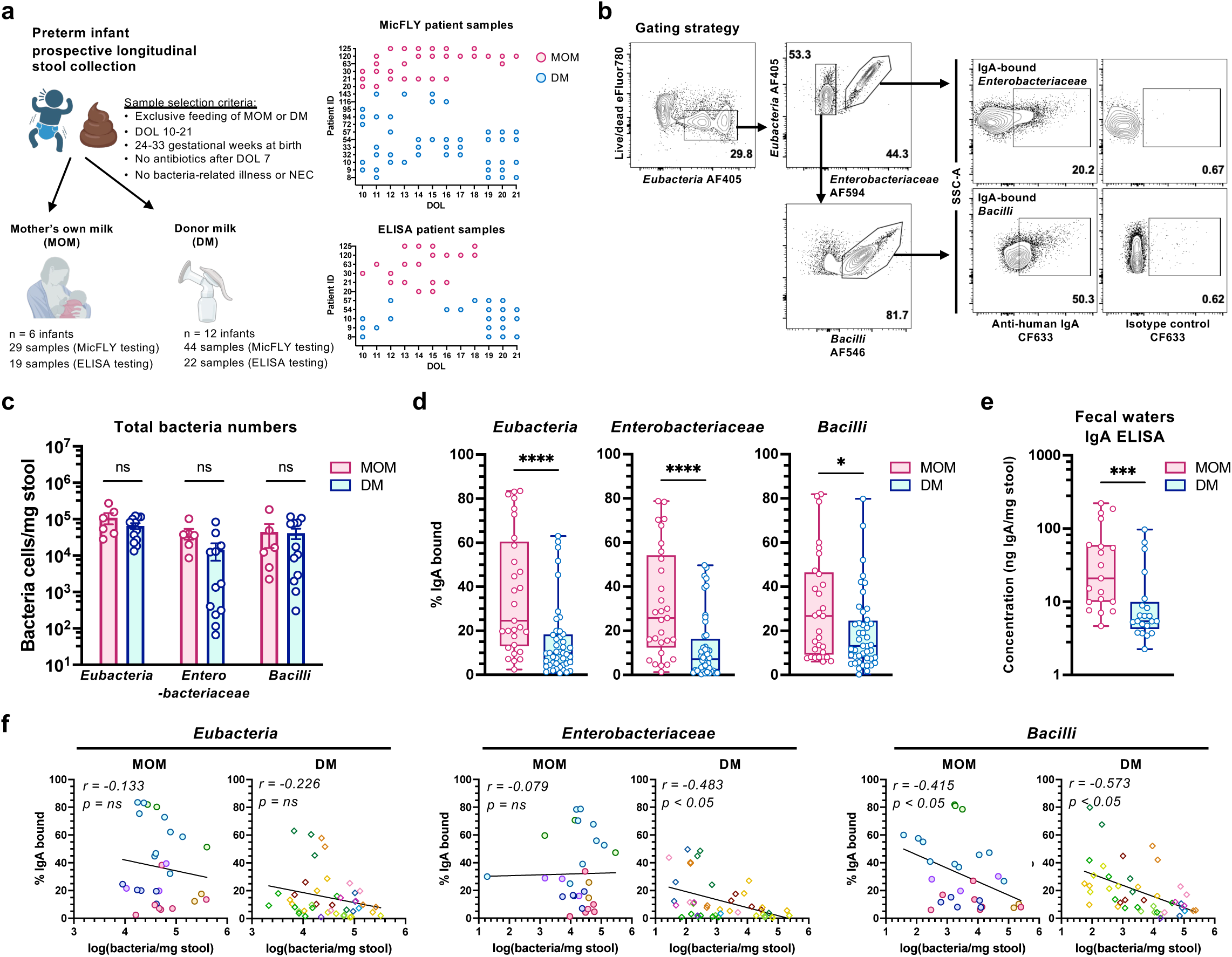
MicFLY reveals differences in IgA binding to microbiota in preterm infants fed mother’s own milk or donor milk. **a**, Study design and sample overview. Graphs summarize longitudinally collected stool. The day-of-life (DOL) on which a sample was collected is shown by an individual dot. Colors indicate feeding group (pink = MOM, blue = DM), and each row corresponds to one infant. **b**, Gating strategy to identify IgA-bound bacteria with MicFLY. IgA+ gating was defined using isotype control antibody (goat anti-chicken IgY CF633). **c**, Absolute abundances of *Eubacteria*, *Enterobacteriaceae*, and *Bacilli* populations measured in MOM- and DM-fed infants and calculated by MicFLY. Bars represent average total counts ± SEM for pooled samples from DOL 10-21 for each infant. Each dot represents an individual infant. Statistical analyses use unpaired two-tailed Welch’s t-test. **d**, MicFLY measurements of percent IgA-bound bacterial taxa compared by feeding type. Aggregated data of all samples is shown. Box plots show median, interquartile range, and minimum to maximum values. Each dot represents a single sample. Statistical analyses use two-tailed unpaired Mann-Whitney test. **e**, ELISA measurements on total concentration of free IgA (ng of IgA per mg stool) compared between feeding type. Aggregated data of all samples is shown. Box plot shows median, interquartile range, and minimum to maximum values. Each dot represents a single sample. Statistical analyses use two-tailed unpaired Mann-Whitney test. **f**, Correlation between bacterial abundance (log-transformed) and percent IgA-bound bacteria within each taxon, measured by MicFLY. Best-fit line from simple linear regression is displayed; *r* and *p*-value from Spearman correlation. Each dot represents a single sample, and each color corresponds to an individual patient. Repeated colors indicate multiple samples per patient. For all statistical tests: p < 0.05 (*), p < 0.01 (**), p < 0.001 (***), p < 0.0001 (****). All analyses done on live cells only.

Because infants begin making their own antibodies around three to four weeks post-delivery^1,89^, we limited analyses to DOL 21 to assess the binding of milk-derived IgA to the microbiota. We measured the two taxa (class: *Bacilli* and family: *Enterobacteriaceae*) comprising the majority of bacteria in neonatal preterm infants (**Figure 4B**)^21,90–92^. Total abundance quantification by MicFLY revealed similar bacterial loads between DM and MOM-fed infants, averaged across DOL 10-21 (**Figure 4C**).

Despite the lack of differences in total numbers, aggregate data revealed a significant decrease in the proportion of IgA-bound *Bacilli* and *Enterobacteriaceae* in DM-fed infants (**Figure 4A**). ELISA measurement of fecal waters confirmed decreased concentration of free IgA in DM-fed infants, supporting that pasteurization of milk reduces infant intestinal IgA levels significantly (**Figure 4E**). We next asked if variations in bacterial density could be affecting intestinal IgA binding. Interestingly, we found that *Enterobacteriaceae* bacterial load had no effect on IgA binding in MOM-fed infants, but higher abundance was correlated with decreased binding in DM-fed (**Figure 4F**). These data indicate that saturation of anti-*Enterobacteriaceae* antibodies may be reached sooner in DM due to reduced concentrations or reactivity^82^. In contrast, for *Bacilli*, increased load significantly correlated with decreased IgA binding in both groups, indicating reactivity differences may also play a role in determining the saturation point for milk-derived antibodies. Together, these findings reveal that alterations in IgA levels due to pasteurization alter antibody binding of bacteria in the intestine of infants fed DM.

### MicFLY identifies that a lack of IgG and IgA binding to *E. coli* is associated with the development of necrotizing enterocolitis

Preterm infants are uniquely at risk for necrotizing enterocolitis (NEC), a deadly intestinal disease with a mortality rate up to 35%^93,94^. NEC is a multifactorial condition culminating in acute intestinal inflammation and necrosis. *Enterobacteriaceae* family bacteria are associated with NEC^1,83^; however, no single species has been identified as the sole cause^90,95^. Breast milk IgA reduces NEC risk^85^, and decreased frequencies of IgA-bound *Enterobacteriaceae* are associated with NEC development^1^. It is not known if this loss in antibody binding over time is due to expansion of certain species not targeted by IgA or whether bacterial overabundance is saturating maternal antibodies in the intestine. Additionally, maternal IgG is becoming increasingly recognized as critical to neonatal immunity and may cooperate with IgA during early life to limit gut bacteria-induced inflammation^96–98^. What role this plays in the preterm microbiota has not been investigated.

To test MicFLY’s ability to quantify the antibody-bound preterm microbiota, we selected longitudinal samples from a small cohort of preterm infants that either did or did not develop NEC. We designed a set of probes enabling detection of the most common taxa found in preterm infants (Class *Bacilli*: *Staphylococcus spp*., *Enterococcus spp*., and *Streptococcus spp*.; Family *Enterobacteriaceae*: *E. coli*, *K. pneumoniae*, *K. aerogenes*, *K. oxytoca*, *E. cloacae*) **(Figure S7A)**^21,83,90,99–101^. This panel permits us to concurrently measure IgG and IgA binding to each taxon **(Figure S7B).** We collected fecal samples from patients with prospective NEC diagnosis (n = 7 infants, 72 samples) or matched control infants who did not develop NEC (n = 6 infants, 71 samples) (**Figure 5A**). Each control infant had at least 1 NEC match based on biological sex, gestational age at birth (+/- 2 weeks), delivery mode (vaginal or C-section) and feeding (MOM/partial MOM or DM exclusively) **(Table S4).** NEC samples were collected prior to diagnosis.

**Figure 5.**
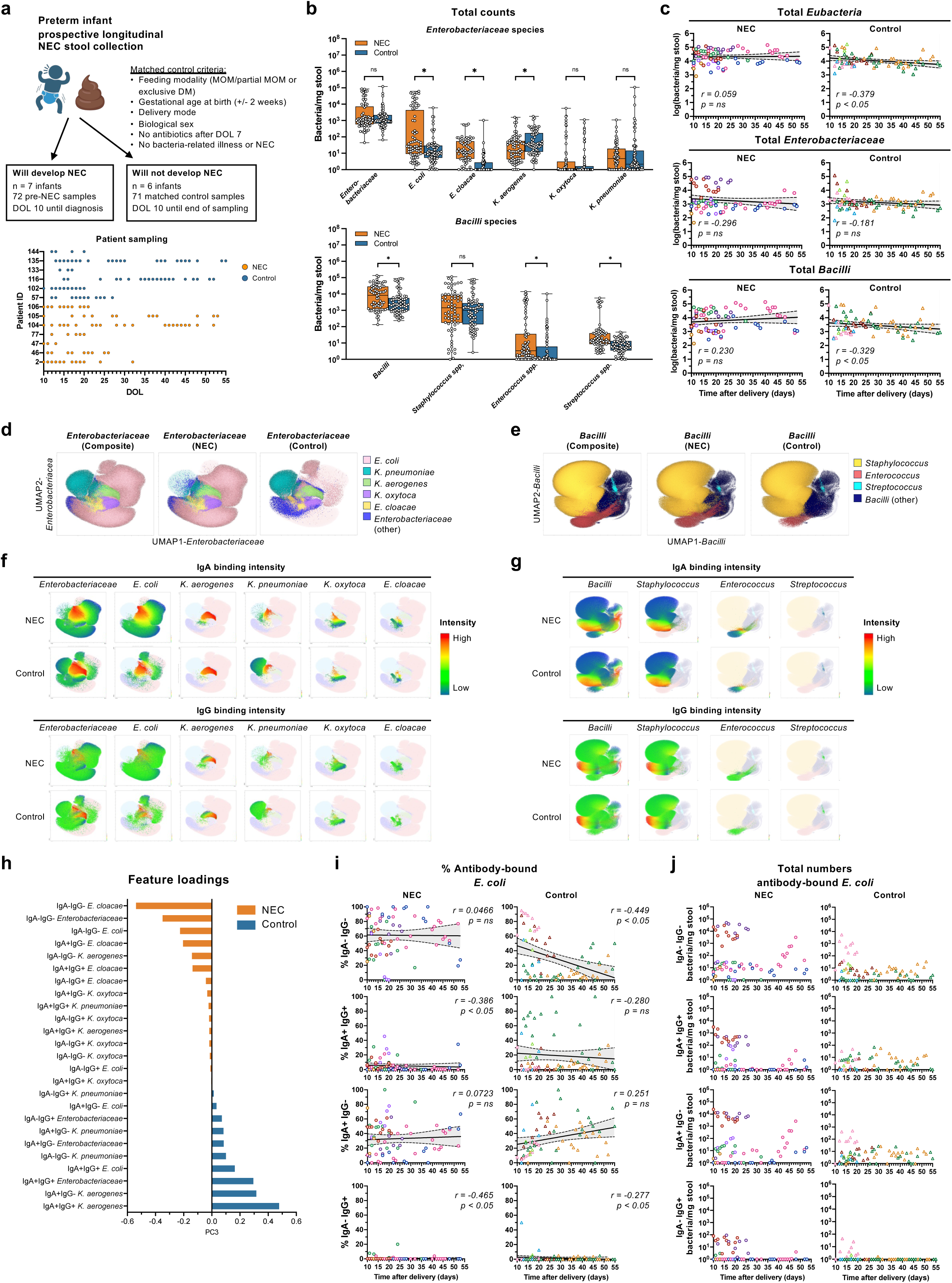
Species-resolved longitudinal antibody binding dynamics in preterm infants who develop NEC. **a**, Study design and sample overview. Graphs summarize longitudinally collected stool. The day-of-life (DOL) on which a sample was collected is shown by an individual dot. Colors indicate patient group (orange = prospective NEC diagnosis, blue = control), and each row corresponds to one infant. **b**, Absolute abundance of *Enterobacteriaceae* and *Bacilli* species measured by MicFLY using genus/species-specific probes. Aggregated data of all samples is shown. Box plots show median, interquartile range, and minimum to maximum values. Statistical comparisons performed using unpaired two-tailed multiple Mann-Whitney tests with Holm-Sidak correction. **c**, Longitudinal correlation of day-of-life (DOL) and total bacterial counts (log-transformed) comparing NEC and control infants for amounts of *Eubacteria*+, *Enterobacteriaceae*+, and *Bacilli*+ bacteria measured by MicFLY. Best-fit line from simple linear regression is displayed, and shaded area represents 95% confidence interval; *r* and *p*-value from Spearman correlation. Each dot represents a single sample, and each color corresponds to an individual patient. Repeated colors indicate multiple samples per patient. **d,** UMAP projections of species-specific *Enterobacteriaceae*+ events as determined by MicFLY. Data was concatenated from all patients and timepoints. UMAPs depict species distribution across NEC and control groups. Plots generated using FlowJo’s UMAP plugin (5 nearest neighbors, minimum distance 0.1, 2 components) and visualized with Cluster Explorer. Each dot represents a single bacterial cell, which is color-coded according to species. **e,** UMAP projections of species-specific *Bacilli*+ events exactly as in panel **d**. **f,** Visualization of IgA or IgG binding intensity overlayed onto UMAP from **d**. Each dot represents a single bacterial cell; color scale reflects anti-IgA or anti-IgG fluorescence intensity, scaled from minimum to maximum within the dataset. Species-specific clusters (*E. coli*, *K. aerogenes*, *K. pneumoniae*, *K. oxytoca*, *E. cloacae*) are shown in separate cluster-level views. **g,** IgA or IgG binding intensity overlays for *Bacilli*+ populations, exactly as in panel **f**. **h**, Sorted feature loadings along Principal Component 3 (PC3) from Joint-Compositional Tensor Factorization (Joint-CTF) analysis using longitudinal absolute abundances of *Enterobacteriaceae* taxa and their antibody binding. Feature rankings for only the IgA/IgG-bound taxa are shown. Features are ordered by their PC3 coefficients, reflecting their relative association with NEC (orange) or control (blue). **i,** Longitudinal correlation of DOL and percent antibody-bound *E. coli* comparing NEC and control patients measured by MicFLY. Antibody binding is split between IgG- and IgA-bound bacteria, as indicated. Best-fit line from simple linear regression is displayed, and shaded area represents 95% confidence interval; *r* and *p*-value from Spearman correlation. Same samples and color coding as in panel **c**. **j**, Longitudinal display of absolute abundance for antibody-bound *E. coli* in NEC and control patients measured by MicFLY. Total numbers are shown for *E. coli* bound by IgA, IgG, or both. Same samples and color coding as in panel **c**. All analyses done on live cells only.

We next used MicFLY to perform quantitative species-level measurements of the microbiota. Aggregated abundance data showed that among the *Enterobacteriaceae*, *E. coli* had the highest load in patients who progressed to NEC and was significantly elevated compared to controls (**Figure 5B**). For *Bacilli* bacteria, the *Staphylococcus* species were the most abundant but did not differ between NEC and control. Low abundance taxa *E. cloacae, Enterococcus* and *Streptococcus* were also higher in NEC, while *K. aerogenes* was enriched in controls. However, we are cautious of drawing conclusions from low abundance taxa at or below the assay’s limit of detection **(Figure S4C).**

We next tested if we could use longitudinal data to associate NEC development to expansion of *Enterobacteriaceae* or *Bacilli* populations. However, supporting previous reports^102,103^, we did not see a correlation between shifts in bacterial load and NEC (**Figure 5C**). To comprehensively visualize our single cell data, we graphed our data as Uniform Manifold Approximations and Projections (UMAPs). UMAP analysis from aggregated data confirmed total abundance findings (**Figure 5B**) that *E. coli* is overrepresented in NEC (**Figure 5D**). In contrast, UMAP analysis of *Bacilli* species produced highly similar patterns for both groups (**Figure 5E**). Day-by-day projections made from each patient **(Figures S8 and S9)** revealed that many NEC-affected infants (Patients 46, 77, 104 and 106), but not controls, are colonized by persistent and/or increasing populations of *Enterobacteriaceae* and in particular, *E. coli*. These results are not distortions produced by UMAP visualization as species-level total abundances calculated for each patient revealed a similar pattern **(Figure S10)**.

We next incorporated antibody binding data into our analyses. UMAPs revealed that clustering of *Enterobacteriaceae* species (**Figure 5F**), but not *Bacilli* (**Figure 5G**), was driven by antibody binding intensity, with several *Enterobacteriaceae* species having high IgA/IgG binding localized to a center cluster (**Figure 5F**). The proportions of antibody-bound and unbound subpopulations also differed between NEC and control (**Figure 5F**), with more IgA/IgG unbound *E.coli* enriched in NEC. Conversely, IgA/IgG unbound *K. pneumoniae* was enriched in the control, but this appeared to be driven by a single patient **(Figure S9).** Quantitative analysis of aggregated longitudinal data revealed significantly reduced IgA and IgG-bound *Enterobacteriaceae* and IgA-bound *Bacilli* in patients who progress to NEC **(Figures S11A and S11B)**, thereby confirming UMAP analyses. IgG also appeared targeted toward the *Enterobacteriaceae*, as there was no significant difference between IgG-bound *Bacilli* between NEC or control (**Figure S11B)**. We asked whether these differences in antibody binding could be due to increased bacteria exceeding intestinal antibody supply, but we observed no significant correlation between higher load and lower binding **(Figure S11C)**. Taken together, these results strongly indicate that deficits in antibody reactivity, not bacterial overabundance, are contributing to observed differences in binding.

To better understand how antibody binding and species-level abundance dynamics might differentiate NEC and control, we utilized a multi-modal dimensionality reduction tool, Joint Compositional Tensor Factorization (Joint-CTF), that takes into account repeated measurements and sparsity in multi-modal microbiome data to identify differentially abundant bacteria throughout the microbiota’s development^104^. Based on our analyses indicating that differences amongst *Enterobacteriaceae* discriminate NEC from control (**Figures 5B, 5D-5G**), we focused on species within that taxon. Joint-CTF analysis showed clear clustering of patients along the third axis of variation (PC3) **(Figure S11D)**, which significantly separated them by diagnosis **(Figure S11E)** and consistently differentiated the two groups over time **(Figure S11F).** The features contributing to the subject-level separation shown in PC3 (i.e., feature loadings) were sorted based on their coefficients, with highest rankings most associated to NEC and lowest rankings associated to control **(Figure S11G)**. The top features associated with NEC were IgA^-^/IgG^-^ bacterial taxa, specifically *E. cloacae, E. coli* and *K. aerogenes* (**Figure 5H**). In contrast, IgA^+^/IgG^+^ abundance for many of these same taxa, including *E. coli,* was associated with control patients (**Figure 5H**).

As confirmation of Joint-CTF, longitudinal correlation analyses of both percent and number of IgA/IgG bound bacteria confirmed nearly undetectable IgA^+^IgG^+^ double-bound *Enterobacteriaceae*, and specifically *E. coli*, in infants who will develop NEC (**Figures 5I, 5J, S11H).** Consistent with IgA^+^IgG^+^ data, the percentage of IgA^-^IgG^-^ *Enterobacteriaceae* and *E. coli* significantly decreases over time in control but not NEC (**Figures 5I and S11H)**. The absolute number of IgA^-^IgG^-^ *Enterobacteriaceae* and *E. coli* was also maintained or increasing in some NEC patients but not controls (**Figures 5J and S11H)**. Interestingly, IgG^+^ bacteria was almost always exclusively IgA^+^, while the converse was not true (**Figures 5I, S7B, and S11H**). These data indicated that lack of IgG binding to *E. coli* was the dominant association with NEC development. The top ranked feature, *E. cloacae* (**Figure 5H**), had too little bacteria to draw meaningful conclusions **(Figure S11I).**

Taken together, our quantitative analyses of the preterm microbiota highlights the extraordinary potential of MicFLY to identify factors influencing microbiota dynamics and proper establishment of the infant gut microbiome. While this cohort is too small to draw actionable conclusions in regard to NEC, we believe our data demonstrates the striking ability of MicFLY to quantify the number and properties of individual species in mixed communities and associate them to disease.

## DISCUSSION

Here we demonstrate MicFLY as a single cell technology that performs absolute quantification of the microbiota using spectral flow cytometry. Various methods have previously been used to quantify the microbiome, which associate relative abundances to total abundances with qPCR^18–20^ or spiked-in sequences^21–24^. However, confounding factors may limit accuracy in these approaches. For instance, loss of signal from low abundance bacteria is common in samples with low biomass, making comparisons between samples unreliable^105,106^. Relative abundances may also be skewed by amplification biases caused by unequal primer/template concentration^25–30,107,108^ or DNA isolation efficiencies^27,31–35^. The number of 16S gene copies also varies by species, causing detection bias toward taxa with higher copy numbers^109,110^. Thus, microbiota abundances have been difficult to compare across laboratories, limiting accurate assessment of the role of differential bacterial abundance in the host^35,111^. MicFly advances beyond these previous methods by enumerating individual bacteria with spectral flow cytometry, thus eliminating sequencing biases. However, because MicFLY is a targeted method, sequencing will still be necessary to uncover unknown microbiotas.

We demonstrate through incorporation of an amine-reactive dye that MicFLY can measure microbiota viability. Most microbiota sequencing methods do not take into account bacterial viability because it is challenging to assess^51,112^. Thus, the impact of bacterial cell death on the gut microbiota and host immunity is not well understood^53–58^.

For example, we know that colonization shifts significantly according to circadian rhythms, but the patterns of bacterial cell life and death are unclear^113–115^. MicFLY’s ability to detect significant quantifiable differences in viability among taxa in the microbiota opens up further investigation into these questions.

Recently, single cell bacterial transcriptomics have been demonstrated on a variety of platforms^116–125^. Unlike eukaryotic single cell RNA-sequencing (scRNA-seq), all bacterial scRNA-seq platforms require extensive pre-processing of the mRNA that potentially introduce bias. In contrast, MicFLY performs single cell mRNA quantification without enzymatic amplification of nucleic acids, permitting direct assessment of transcripts. A substantial breakthrough of MicFLY is the combination of mRNA- and taxon-targeted probes to measure bacteria-specific gene expression within mixed populations. This aspect of MicFLY has a plethora of applications, such as investigating how heterogeneous bacterial populations drive host responses during infection and chronic inflammatory disease.

By incorporating surface protein-targeted fluorescent antibodies, we demonstrate MicFLY can quantify the antibody-bound microbiota. Previously, antibody-bound bacteria could only be studied using Ig-Seq, a method requiring comparison of sorted populations that is limited to non-quantitative, compositional analyses. Ig-Seq can only determine whether an antibody-bound population is enriched with a given taxon, but not how much each taxon is directly bound by antibodies^6^. In contrast, MicFLY directly measures binding at the single cell level with concurrent IgA/IgG measurements, revealing independent and overlapping antibody-bound bacteria, as well as preferential antibody targeting at the species-level. MicFLY also improves on Ig-Seq in that it functions robustly in low biomass samples (such as preterm infant fecal samples), whereas species level discrimination with Ig-Seq requires much more material^11^.

To demonstrate the effectiveness of MicFLY, we analyzed stool samples from a prospective longitudinal cohort of preterm infants that did or did not develop NEC. Critically, we demonstrated that eight individual taxa from two families could be quantified at the genus/species level alongside IgA and IgG binding. Even in this small cohort, MicFLY analysis provides highly granular information on longitudinal shifts in the microbiota and their relationship to disease. We previously identified that infants who develop NEC were enriched in IgA unbound bacteria^1^, but here, using better technology, we show an additional important role for IgG. Indeed, even in aggregated longitudinal data, IgG binding of *Enterobacteriaceae* is distinctly enriched in control patients, which we speculate are controlling pro-inflammatory bacteria during early life^96–98^. We hypothesize that a larger cohort could elucidate how IgG and IgA may serve different and/or complementary functions influencing the development of the preterm infant microbiota in both health and disease.

A potential confounder to our NEC cohort data is the presence of DM-fed infants, which we demonstrate have much less IgA bound to intestinal bacteria before DOL 21.

However, we would point out that samples from all DM-fed infants in our NEC study (2/7 of NEC and 1/6 of control) were collected for over 50 days, but NEC-affected infants are dominated by IgG/IgA unbound bacteria long after they should be producing their own antibodies (around 3-4 weeks of life)^1,89^. These findings beg the question of why only some infants fail to produce sufficient amounts of microbiota-reactive antibodies.

Another confounder is that four infants in the NEC cohort, but none in the control, received antibiotics after DOL 7. However, antibiotics were given either far before (patients 104 and 105) or just before (patient 77) diagnosis, thus, we think is not critical to the associations identified by MicFLY. A fourth infant (patient 47) also provides just three samples, which is of limited value overall. Despite these limitations, our work still displays the incredibly high resolution analyses that MicFLY can provide to elucidate microbial signatures of disease; a larger cohort would therefore be needed to precisely define colonization dynamics of the NEC microbiota.

Taken together, MicFLY has the potential to revolutionize microbiome research by enabling quantitative single cell analysis of mixed bacterial populations without the need for sequencing or culturing. MicFLY is a highly versatile technology that provides granular information on bacterial identity, viability, transcription and antibody binding. As such, MicFLY holds extraordinary promise as a tool for future research requiring highly dimensional quantitative data to uncover the dynamics driving microbiome-related disease.

## Supporting information

CLC Files

Probe Binding sites

## ACKNOWLEDGMENTS

This work was supported by the National Institutes of Health (TH: R01CA269902, R01DK120697; CT: F30HD114316; WD R01AI170607; MS 5R01DK133453; LS DP2AI185753;) the Kenneth Rainin Foundation (TH), the Burroughs Wellcome Foundation (TH) and the UPMC Children’s Hospital of Pittsburgh Foundation. We would like to thank Y. Belkaid (NIH/NIAID) for providing IgA -/- mice. We thank Manuella Raffatellu (UCSD) and Leigh Knodler (Univ. of Vermont) for *Salmonella* strains used in the study. We thank E. Hartigan, M. Butoryak, L. Shaver, E. Cribbs, M. McDermott and the staff at the UPMC Magee Women’s Hospital for consenting and collecting infant fecal samples for this study. We thank members of the Hand, Poholek, Silverman, DePas and Shenhav labs for helpful discussions. We thank J. Michel for maintaining the Rangos Research Flow Core and DeWayne Falkner for assisting with cell sorting. We thank Y. Wang and G. Church (Harvard University) for assistance with hairpin design.

We thank the staff of the Division of Laboratory Animal Services for animal husbandry and the Center for Biological Imaging (S. Watkins) for training and use of the confocal microscope used in this study. Finally, we would like to thank the patients and their families for their generous donations of samples to the study.

## AUTHOR CONTRIBUTIONS

Conceptualization, C.M.T., D.A.A., W.H.D., T.W.H.; Data curation, C.M.T.; Formal analysis, C.M.T., B.C.V.; Funding acquisition, C.M.T., M.A.S., L.S., W.H.D., T.W.H.; Investigation, C.M.T., A.R.L., T.C.T., I.A.V., S.W., V.L., K.J.S.; Methodology, C.M.T., B.C., D.A.A., M.A.S., L.H., W.H.D., T.W.H.; Project administration, C.M.T., M.B., T.W.H.; Resources, M.B., M.A.S., L.S., W.H.D., T.W.H.; Supervision, C.M.T., M.A.S., L.S., T.W.H.; Validation, C.M.T., B.C.V., A.R.L, T.C.T., I.A.V., S.W., V.L.; Visualization, C.M.T., B.C.V.; Writing – original draft, C.M.T., B.C.V., M.A.S., L.H., W.H.D., T.W.H.; Writing – review & editing, C.M.T., B.C.V., M.A.S., L.H., W.H.D., T.W.H.

## RESOURCE AVAILABILITY

### Lead contact

Requests for further information and resources should be directed to and will be fulfilled by the lead contact, Dr. Timothy Hand.

### Materials availability

All unique/stable reagents generated in this study are available from the lead contact with a completed materials transfer agreement.

### Data and code availability

- Raw data generated from 16S rRNA gene amplicon sequencing for PedsCom dataset in this study are available as FASTQ files and have been deposited to FigShare (https://figshare.com/s/93ad90f14bf95e046f26).
- This paper does not report original code.
- Any additional information required to reanalyze the data reported in this paper is available from the lead contact upon request.

## EXPERIMENTAL MODEL AND SUBJECT DETAILS

### Preterm infant fecal samples

The human study protocol was approved by the Institutional Review Board (STUDY19110318) of the University of Pittsburgh. This work was conducted as part of a prospective longitudinal cohort study of preterm infants hospitalized in the NICU at UPMC Magee-Women’s Hospital. Written informed consent was obtained from parents/guardians prior to enrollment. Enrolled infants were born between 24 and 33 weeks gestational age with no other exclusions. Fecal sample collection began at day of life (DOL) 10 and ended within 63 days post-delivery or when the infant reaches 33 weeks gestational age, whichever comes first. Sample collection stops when infant is diagnosed with NEC. Fecal samples were collected from the diaper and immediately frozen at −80°C. Samples from all infants who developed NEC were included in this study regardless of antibiotic status due to the limited number of diagnosed patients.

Except for infants who developed NEC, samples selected for testing were only from infants who did not develop any other bacteria-related illness and did not receive any antibiotics after DOL 7.

### Mice

C57BL/6 and 129X1/SvJ mice were purchased from Jackson Laboratories (Bar Harbor, ME). Igha^−/−^ mice were obtained from Yasmine Belkaid (NIH, Bethesda, MD). Non-gnotobiotic mice were maintained under specific pathogen-free conditions. Germ-free C57BL/6 mice were maintained at University of Pittsburgh Gnotobiotic Core facility.

Germ-free and gnotobiotic Eα16/NOD PedsCom colonized mice were maintained at the Hill Pavilion gnotobiotic mouse facility, University of Pennsylvania. NOD mice were colonized with the PedsCom consortium^60^ by cohousing germ-free female NOD mice with PedsCom-colonized C57/BL6 female mouse for 2 weeks. Colonization of cohoused PedsCom NOD mice was confirmed using qPCR probes for each PedsCom taxa as previously described^60^. All germ-free and gnotobiotic mice were maintained under strict germ-free conditions. Mice at Hill Pavillion were housed in flexible film isolators (Class Biologically Clean, WI, USA), fed autoclaved LabDiet 5021 (Cat# 0006540) ad libitum, and caged on autoclaved Beta-chip hardwood bedding (Nepco, NY, USA). Sterility checks were regularly performed on isolators each month and additionally prior to any transfer of animals. Freshly collected pellets were cultured on brain heart infusion (BHI) (Oxoid, UK), NB1, and Sabouraud media for 65–70 hours at 37°C under aerobic and anaerobic conditions with positive and negative control samples. All mice were housed in American Association for the Accreditation of Laboratory Animal Care (AAALAC)-accredited facilities at the University of Pittsburgh or at the University of Pennsylvania.

All animal procedures were conducted in accordance with the Guide for the Care and Use of Laboratory Animals under protocols approved by the Institutional Animal Care and Use Committees (IACUC) at the University of Pittsburgh and University of Pennsylvania.

### *Salmonella* infection model

129X1/SvJ mice, aged 8-16 weeks, were orally pretreated with streptomycin 20 mg. The following day, mice were orally infected with 2.5 × 10^8 colony-forming units (CFU) of *Salmonella enterica serovar Typhimurium* (Strain SL1344)^81^ diluted in PBS. Mice were sacrificed at four hours post-infection and four days post-infection, and cecal and colon contents were collected at both time points.

## METHOD DETAILS

### Preterm infant stool microbiota extraction

Approximately 20 to 200 mg of stool is weighed and recorded. Fecal material is homogenized by resuspending with vigorous vortexing and pipetting in 1 mL phosphate buffered saline (PBS, pH 7.4) then pelleted by centrifugation (10 minutes, 9000g).

Supernatant is collected and stored (−80°C) for testing fecal waters. To remove undigested milk protein/fat, the fecal pellet is washed twice in 2 mL of 5% Percoll/0.15 M NaCl, pelleted (10 minutes, 10,000g) and resuspended in 1 mL PBS. To separate eukaryotic and prokaryotic cells, the sample is centrifuged (5 minutes, 50 g), filtered (20 μm cell strainer; Fisher Scientific #NC9912653) and pelleted (10 minutes, 9000g). If the sample is not immediately to be processed for MicFLY, it is resuspended in 1 mL of 50% glycerol (diluted in molecular biology grade water), aliquoted into replicate tubes with 200-500 uL each, and frozen (−80°C). Final volume of resuspended microbiota (1000 uL in 50% glycerol) is recorded for absolute abundance calculations.

### Mouse stool and intestinal microbiota extraction

Microbiota extraction for mouse stool is similar to preterm infant stool but simplified because mouse fecal matter contain less undigested material. Thus, the Percoll washing steps can be omitted, and stool homogenate can directly proceed to the low-speed step first (5 minutes, 50 g) before continuing protocol exactly as for preterm stool. Microbiota extraction from mouse cecal and colonic contents from the *Salmonella* infection model is exactly as for mouse stool.

### MicFLY protocol

MicFLY can be performed in either 96-well plates or microcentrifuge tubes. The protocol for 96-well plates is for microbiota extracted from stool. The protocol is adapted to microcentrifuge tubes if extracting bacteria from whole cecal or colonic contents. PBS is used at pH 7.4 unless otherwise indicated as alkaline PBS, which is pH 8.0.

#### Viability staining

Aliquoted extracted microbiota stored at −80°C is removed to thaw. For a single test on preterm infant microbiota, 200 uL of sample is pipetted to a new microcentrifuge tube with 1 mL PBS. For a single test on mouse microbiota, 50 uL of sample is transferred. If several tests are needed (i.e., heat-killed vs. non heat-killed samples or for single color controls), several tubes are prepared. To calculate absolute abundances, the volume of each sample is recorded. Samples are pelleted (10 minutes, 10,000g), resuspended in 200 uL PBS and transferred to a 96-well U-bottom polypropylene plate (Greinier Bio-One #650201). For heat-killed mouse stool experiments, prior to transferring to multiwell plate, the resuspended sample was heated at 95°C on a dry heat block for 20 minutes then transferred to plate. Sample is pelleted in the 96-well plate (10 minutes, 4200g) and resuspended with 100 μL fixable viability dye eFluor780 (1:1000 in PBS; Thermo Fisher Scientific #65-0865-18) and incubated on ice in the dark for 30 minutes. Sample is washed with PBS and ready for surface protein staining. If fluorescent antibodies will not be added, sample proceeds directly to PFA fixation step.

#### Fluorescent antibody staining and cross-linking

Fluorescent antibodies are added and incubated on ice for 1 hour in the dark. For IgA binding tests in preterm infants, goat anti-human IgA (alpha chain) polyclonal antibody CF633 (Biotium #20427) or mouse anti-human IgA monoclonal antibody VioGreen (Miltenyi clone IS11-8E10) are used at 1:10 dilution. For isotype controls, goat anti-chicken IgY antibody CF633 (Biotium # 20126) or mouse IgG1 isotype control antibody VioGreen (Miltenyi clone IS5-21F5; #130-113-207) was used at 1:10 dilution. For preterm infant IgG binding tests, mouse anti-human IgG monoclonal antibody BUV737 (BD BioSciences clone G18-145) is used at 1:20 dilution. Antibodies are diluted in 10% normal goat or normal mouse serum in PBS, or both if combining anti-IgA and anti-IgG antibodies. For mouse IgA binding tests, goat anti-mouse IgA (alpha chain) polyclonal antibody DyLight488 (Abcam #ab97011) is used at 1:10 dilution with 10% normal goat serum in PBS. After incubation, sample is washed three times with alkaline PBS (pH 8.0), which is necessary for downstream BS3 crosslinking. After washing, sample is resuspended with 50 μL of PBS at pH 8.0 and mixed with 50 μL of 2 mM BS3 (diluted in alkaline PBS; final concentration = 1 mM) to crosslink antibodies to the bacterial cell surface. Sample is incubated on ice in the dark for 30 minutes. After incubation, BS3 is washed from sample with PBS.

#### Bacterial paraformaldehyde fixation

Sample is resuspended in 100 μL of PBS then mixed with 100 μL of 8% PFA (final concentration 4% PFA). Sample is incubated on ice in the dark for 30 minutes, then washed with PBS.

#### Bacterial permeabilization

To permeabilize bacteria prior to adding initiator probes, fixed bacteria are incubated at 37°C for one hour with 100 μL of 0.1 mg/mL lysozyme (Sigma Aldrich #L6876-1G) and 0.1 mg/mL lysostaphin (Sigma Aldrich #L7386-15MG) diluted in 10 mM Tris-HCl pH 7.0. For mRNA detection, 0.5 units/μL NxGen RNAse inhibitor (BioSearch Technologies #30281-1) is also added to permeabilization buffer, and proper precautions are taken using RNAse-free reagents or DEPC-treated water to dilute buffers going forward. After lysozyme/lysostaphin incubation, 100 uL of PBS with 0.1 % Tween (PBS-T) is added. Sample is pelleted (10 minutes, 4200g), then washed again with PBS-T.

#### Hybridization Chain Reaction (HCR)

For information on probe design see section “Ribosomal RNA database and rRNA initiator probe design”. Prior to adding initiator probes, sample is resuspended with 100 μL pre-warmed probe hybridization buffer (Molecular Instruments) for a minimum of one hour (37°C). While sample is pre-hybridizing, the probe solution containing the panel of initiator probes is prepared. For a single test, 1 µL of initiator probe from 4 µM working stock solution is used. If competitor probes are needed, then 1 µL of a competitor probe from a 20 µM working stock solution is used. All initiator probes in the probe panel (1 μL each probe) are pooled together into 50 μL of probe hybridization buffer per sample.

After the pre-hybridization incubation step, 50 μL of pooled initiator probes in probe hybridization buffer is added to each sample and incubated overnight (37°C, covered). For single color controls, *Eubacteria* initiator probes are used as a positive control to confirm signal detection.

After hybridization, the sample is washed three times with 200μL probe wash buffer (Molecular Instruments; 10 minutes, 4200g) with 5 minute 37°C incubations between washes. Sample is resuspended with 25 μL pre-warmed (room temperature) amplification buffer (Molecular Instruments) to prepare for amplification (30 minutes, room temperature). During incubation, fluorescent hairpins are prepared. 0.35 μL/sample of each fluorescent hairpin (stored at −20°C as a 12 μM solution in molecular biology grade water) is snap-cooled in PCR strip tubes in a thermocycler (95°C for 90 seconds, then 22°C for 30 minutes) then pooled into 15 μL amplification buffer per sample. When pre-amplification is complete, pooled hairpins are added and incubated for 4-18 hours (room temperature; covered)

After hairpin amplification, samples are washed with 200 μL 5X SSCT (saline sodium citrate with 0.1% Tween) three times, (10 minutes, 4200g) and resuspended in a final volume of 200 μL. This final volume must be recorded for absolute abundance calculations.

#### Spectral flow cytometry

Spectral flow cytometry was performed on the Cytek Aurora (Cytek Biosciences). Scatter gain settings were: FSC-A = 250, SSC-A = 50, and SSC-B = 50. Threshold settings used the “AND” function, set to 1500 for FSC-A, 1500 for SSC-A, and between 500-1000 for the peak fluorescence channel corresponding to the *Eubacteria* probe/hairpin pair used in the spectral panel. For example, when *Eubacteria* probe with an Alexa Fluor 405-conjugated hairpin was used, a threshold was set on the V2 channel. Stopping gate is placed on at least 20,000 live *Eubacteria*+ events. To analyze rare events, 100,000 live *Eubacteria*+ events are captured. Some preterm infant stool had very few bacteria with a minimum of 5000 live *Eubacteria*+ events captured. Cytek volumetric counting is automatic and obtained using $VOL function on FlowJo v10.10.

### Bacterial culture

For single culture experiments using multiplexed probe panels to target all major gut bacterial taxa, facultative and obligate anaerobic species were grown under a range of conditions, detailed in **Supplemental Table 4**. All facultative anaerobic bacteria, except *L. gasseri,* were streaked from −80°C stocks (25-50% glycerol in LB media) onto LB, MacConkey, or tryptic soy agar (TSA) plates and incubated at 37°C overnight prior to growing in liquid culture. A single isolate was then inoculated into 5 mL LB broth and incubated shaking at 37°C for ∼18 hours. *L. gasseri* was directly seeded into 5 mL MRS broth from frozen stock and incubated shaking at 37°C for 48 hours. For obligate anaerobic bacteria, cultures were grown in an anaerobic chamber (0 ppm O_2_, 2.0% H_2_). Anaerobic bacteria were streaked from −80°C glycerol stocks onto TSA with 5% sheep blood or modified clostridial agar and incubated at 37°C for 24-72 hours. A single isolate was used to inoculate 2-5 mL broth (reinforced clostridial medium or peptone yeast extract broth; Bifidus Selective Medium Broth for *Bifidobacterium* species) and incubated for another 24-96 hours (**Table S5**). For some strains in original manufacturer vial, bacteria were either fixed directly or inoculated into broth immediately. For PedsCom isolates, bacteria were grown as previously described^60^. The volume of bacteria culture used for MicFLY testing was the amount needed to obtain a visible pellet which differed depending on species and was sample-specific (∼20-200 μL of overnight culture for facultative anaerobes; ∼100-1000 μL for obligate anaerobes). Prior to MicFLY testing, all cultures are washed with PBS, and fixed with 4% PFA.

For mixed culture experiments, bacteria were grown separately and combined after fixation. For *E. coli*, *S. aureus*, and *B. breve* mixed culture experiments, 25 μL from overnight *E. coli* and *S. aureus* cultures were washed, fixed, and combined with 250 μL of fixed *B. breve* from a 1 mL culture. For *E. coli* and *Stm* mixed culture experiments, 4 x 10^7^ CFU of each bacteria were used.

### Reporter GFP *E. coli* and mCherry Stm cultures

For GFP mRNA detection experiments, DH5α *E. coli* strain with arabinose-inducible GFP carried on p006-GFP-pBAD plasmid with a kanamycin resistance gene (Addgene #108315) is streaked onto kanamycin agar plates (50 μg/mL) and incubated overnight at 37°C. An isolate is inoculated into 5 mL kanamycin LB broth (100 μg/mL) and incubated shaking overnight at 37°C. A 1:100 dilution of overnight culture is seeded into fresh kanamycin LB broth the next day, and bacteria are grown to log phase (3-4 hours at 37°C) prior to arabinose induction. For dose-response experiments, different *E. coli* cultures were induced for three hours with arabinose final concentrations of 0%, 0.0001%, 0.0005%, 0.001%, 0.005%, 0.01%, 0.05%, or 0.1% (w/v). As a control, non-transformed *E. coli* (605 strain^82^) is grown and tested in parallel in kanamycin-free media. For time-course experiments, two *E. coli* cultures were grown and induced with either 0.01% or 0% arabinose. A fraction of each culture was assayed at defined intervals post-induction (0, 15, 30, 45, 60, 120, 180, and 240 min). In induction experiments with spent media, three cultures of GFP *E. coli* were induced with 0.01% arabinose for 1 hour, 4 hour, or 18 hours. At each time point, bacteria were pelleted (5 minutes, 9000g), and ‘spent’ media was collected, filtered through a 0.2 micron syringe filter and stored at 4°C. Three new GFP *E. coli* isolates were grown, and overnight bacteria were used to seed a fresh culture grown to log phase. Spent media from the different time points were used to induce *E. coli* for one hour.

For mCherry mRNA detection, *Stm* (wildtype or mCherry-expressing strain) is streaked onto MacConkey plates and incubated overnight at 37°C. An isolate is inoculated into 5 mL LB broth and incubated shaking overnight at 37°C. 250 μL of overnight culture is seeded into 5 mL of fresh LB broth the next day, and bacteria are grown to log phase (3-4 hours at 37°C) prior to assaying transcripts. 100 μL of log phase culture is used for MicFLY testing in these experiments. To prevent mRNA degradation, isolated bacteria are immediately fixed (4% PFA) and tested.

### GFP sorting and quantification

To quantify the 16S or ompX gene from GFP-expressing *E. coli*, *E. coli* were induced with 0.01% arabinose for one hour as described above. For sorting, 200 uL of induced bacteria are washed and 4% PFA-fixed, then undergo the MicFLY protocol using a panel with *Eub338* probe/B3-Alexa Fluor546, *Enterobacteriaceae* probe/B2 Alexa-Fluor 546, and GFP mRNA probe/B1-Alexa Fluor 647. MicFLY-treated bacteria are resuspended in 500 uL of 5X SSCT in FACS tubes and sorted on the Cytek Aurora Cell Sorter (threshold on FSC-A = 500). 2,000,000 events, or as many as possible until sample ran out, were sorted separately for Eub338+GFP+ and Eub338-GFP-populations, and bacteria are pelleted (5 minutes, 9000g). To extract DNA from sorted cells, nucleic acids are first decrosslinked by resuspending pellet with lysis buffer (50 mM Tris pH 8.0, 200 mM NaCl, 25 mM EDTA) and proteinase K (0.4 ug/uL; BioSearch technologies #MPRK092) then incubated at 60°C for 1 hr. QIAamp PowerFecal Pro DNA Kit (Qiagen) was then used according to manufacturer’s instructions, with a minor modification in the initial lysis step: 500 µL of solution CD1 was added to each sample, vortexed for 15 seconds, and centrifuged (1 minute, 5000g) before proceeding with the standard protocol. Extracted DNA is stored at −20°C.

To quantify bacterial genes by qPCR, input DNA was normalized by using a volume corresponding to 200,000 sorted events. SsoAdvanced Universal SYBR Green Supermix (BioRad #1725271) was used according to manufacturer’s instructions.

Briefly, 5 uL SYBR green, 1 uL forward primer (20 uM), and 1 uL reverse primer (20 uM) were mixed with diluted DNA in nuclease-free H2O up to a 10 uL reaction. Forward primer for ompX gene was 5’-GCTTTCACCGCAGGTACTTC-3’. Reverse primer for ompX gene was 5’-ATGCTTGCCCAGTCGTTAAT-3’. Forward primer for 16S gene was 5’-GTGCCAGCMGCCGCCGTAA-3’. Reverse primer for 16S gene was 5’-CCGYCAATTYMTTTRAGTTT-3’ Data acquisition was performed on the CFX Duet Real-Time PCR System (Bio-Rad).

### MicFLY vs. Syto BC quantification

To compare MicFLY and Syto BC methods to colony-forming units (CFUs), isolates of *E. coli*, *K. pneumoniae*, *S. aureus*, and *E. faecalis* are grown as described. To serially dilute bacteria, 100 μL of overnight culture is diluted into 900 μL LB broth (dilution factor 10^1^), and repeated for each previous dilution until dilution factor 10^10^ is reached. To count CFUs, 100 μL of each dilution is plated onto MacConkey agar (*E. coli* and *K. pneumoniae*) or LB agar (*S. aureus* and *E. faecalis*) in duplicate (thus increasing the dilution factor another tenfold), followed by overnight incubation at 37°C. CFUs are calculated the next day. The remaining 900 uL of diluted sample is pelleted (5 minutes, 9000g), washed with PBS, fixed with 4% PFA on ice for 30 minutes and resuspended in 900 μL PBS. 100 μL from each dilution tube is set aside at 4°C for Syto BC testing. A 100 μL is aliquot is then assayed with MicFLY. The other fraction is then incubated with Syto BC (Thermo Fisher #S34855) at 1:400 dilution for 15 minutes on ice, washed 3 times with PBS, and resuspended in 100 μL PBS. Samples are run in parallel according to our MicFLY spectral cytometry settings. Counts from colony growth are then compared to results from the flow cytometer.

### Bacterial imaging

To image bacterial cultures, 20 μL of an overnight *E. coli* culture and 200 μL of an overnight *S. aureus* culture are fixed, mixed together, and undergo HCR according to the MicFLY protocol using a panel with *Enterobacteriaceae* probe/B2-Alexa Fluor 594 and *Bacilli* probe/B4-Alexa Fluor 488. After MicFLY treatment, bacteria are pelleted (5 minutes, 9000g) and resuspended in 5 μL of PBS. The 5 μL bacterial suspensions are then mixed with 20 μL of 2% low melting point agarose (Thermo Fisher #16520050), and 10 μL is loaded onto individual wells of a 10-chamber imaging spacer (GraceBio #470352) on a glass microscope slide. Bacteria are imaged on a Leica TCS SP8 confocal microscope using 60x objective.

### *In vitro* breast milk IgA experiment

*In vitro* IgA binding to bacteria was performed according to our previous protocol^82^. Briefly, 30 μL of an overnight *E. coli* culture (strain #605) was incubated with or without IgA purified from breast milk in 10% normal goat serum on ice for 1 hour. After incubation, bacteria was washed 3 times in PBS then stained with secondary antibody goat anti-human IgA (alpha chain) polyclonal antibody CF633 (Biotium #20427) in 10% normal goat serum in PBS on ice for 1 hour. Bacteria were washed 3 times with alkaline PBS (pH 8.0) to prepare for antibody crosslinking/fixation exactly according to the 1 mM BS3 and 4% PFA steps of the MicFLY protocol. After PFA fixation, sample was fractionated into two tubes, in which one proceeds through the rest of the MicFLY protocol with probe hybridization and HCR, while the other is stored at 4°C. Prior to running on the spectral flow cytometer, both fractions are stained with Syto BC (Thermo Fisher #S34855) at 1:400 dilution for 15 minutes on ice then washed 3 times with PBS. Samples are run according to our MicFLY spectral cytometry settings.

### Fecal waters IgA ELISA

Fecal waters were prepared using 1 mL of preterm infant stool homogenate in PBS as described (see ‘**Preterm infant stool microbiota extraction**’). Human IgA was quantified using a commercial ELISA kit (Abcam #ab137980) according to manufacturer’s protocol. Because fecal IgA concentration is unknown, fecal waters were tested at 1:20 or 1:50 dilutions. Samples for each dilution were assayed in duplicate wells and averaged. OD was read at 450 nm on a microplate reader (BioTek Synergy H1). To normalize data, IgA concentration (ng) in 1 mL fecal waters was divided by original stool mass and displayed as ng IgA/mg stool.

### DNA Extraction for PedsCom qPCR and 16S-seq

A stool pellet from each PedsCom mouse was collected into a microcentrifuge tube and weighed prior to DNA extraction. DNA was isolated using QIAamp PowerFecal Pro DNA Kit (Qiagen #51804) according to manufacturer’s instructions with the following modification: DNA was eluted in 50 µL of Solution C6 after heating column at 56°C for ∼10 min to increase yield. DNA was also extracted from germ-free mouse stool as a negative control. Eluted DNA was stored at −20°C. qPCR and 16S sequencing were tested using the same DNA sample.

### qPCR absolute quantification for PedsCom mice

TaqMan custom primer and probe sets were designed to target the DNA-directed RNA polymerase subunit beta (rpoB) gene for each PedsCom member as previously described^60^. Primer and probe sets were generated using the PrimerQuest Tool (Integrated DNA Technologies, IA, USA) and compared with the Multiple Primer Analyzer tool (Thermo Fisher). Absolute quantification of each PedsCom member was determined by rpoB gene copies per gram feces. Data acquisition was performed on the CFX Opus 96 PCR system using TaqMan Multiplex master mix (Thermo Fisher). Data analysis was performed on CFX Maestro ver. 2.3 (Bio-Rad).

### 16S rRNA gene amplicon sequencing and analysis

Extracted DNA from PedsCom or germ-free stool was sent to Microbiome Insights (Vancouver, BC, Canada) for 16S gene amplicon sequencing (V4 region) using 515F/806R primers with Illumina adapters. Library preparation followed the Illumina 16S Metagenomic Sequencing Library Preparation protocol, and paired-end sequencing (2 × 250 bp) was performed on an Illumina MiSeq instrument. Raw demultiplexed FASTQ files were returned for downstream analysis.

Sequencing analyses were conducted in QIIME 2 (version 2024.5) with default parameters unless specified. Demultiplexed paired-end reads were imported via a PairedEndFastqManifestPhred33V2 manifest file, and quality profiles were inspected for trimming/truncation parameters. Only forward reads were retained for denoising because reverse reads exhibited reduced quality and insufficient overlap for merging.

The first five bases of each forward read were trimmed with truncation at 158 bp to remove low-quality tails. DADA2 denoising applied quality filtering, dereplication and consensus-based chimera removal, producing an amplicon sequence variant (ASV) table and representative sequences. Taxonomic assignment was performed with the feature-classifier plugin (classify-sklearn) using a pre-trained naive Bayes classifier trained on the 515F/806R V4 region of the SILVA 138 (SSURef NR99) database (QIIME 2 release 2022.10; scikit-learn v1.4.2). Only samples >10,000 reads (post-denoising) were included with Archaea features removed. ASV counts were normalized to total reads per sample to calculate relative abundance, which was visualized with the QIIME 2 taxa barplot tool. Relative abundance outputs were exported and plotted in GraphPad Prism 10.

### Ribosomal RNA database and rRNA initiator probe design

A reference database of bacterial rRNA sequences was created using CLC Main Workbench v8.1 (Qiagen). For 16S rRNA, all sequences were downloaded as a single FASTA file from BioProject 33175 (Accession: PRJNA33175) from the NCBI RefSeq Targeted Loci Project, which was imported into CLC Main Workbench. For 23S rRNA, BioProject 188943 (Accession: PRJNA188943) was used. At the time of our database generation, 26020 16S sequences and 1350 23S sequences were available for download. Sequence lists were created on CLC Main Workbench to organize 16S genes manually into folders for each taxonomic rank of bacteria relevant to our study, classified based on the List of Prokaryotic names with Standing in Nomenclature (LPSN) which complies with The International Code of Nomenclature of Prokaryotes (ICNP). These organized 16S files are available for download in the **Data S1**. Though not all bacteria were organized into taxonomic categories in our database, we included the ones most relevant to our studies and the human gut microbiota.

To design rRNA initiator probes, CLC is first used to perform gene alignments on 16S or 23S rRNA for target taxa along with a number of closely related “off-target” bacteria.

Several 16-24 nucleotide long regions of interest that perfectly match target taxa, but not off-target taxa, are selected as ‘candidate probes’ to cross-reference against the rRNA database. The “Find binding sites and create fragments” tool is used, the corresponding rRNA database is selected, and each candidate probe is tested separately, limiting maximum mismatches displayed to 2. To select a probe for MicFLY testing, the probe must have 0 mismatches to target bacteria only. Probes with 0 mismatches to off-target bacteria were discarded for further evaluation. Additionally, there must be a limited number of off-target bacteria with 1 mismatch to the candidate probe, such that competitor probes can feasibly be generated to block all possible mismatches on off-target bacteria. Competitor probes were generated by creating sequences that have 0 mismatches to off-target bacteria and pooled with initiator probes during the hybridization step at 5x concentration (20 μM working stock concentration for competitor probes; 4 μM working stock concentration for initiator probes). From empirical observations, 2 mismatches to off-target bacteria do not result in detectable signal from probe hybridization. If a candidate probe has mismatches with several target bacteria, additional probes can be designed to expand coverage at that site, thus making a “probe set” (**Table S1**). Once a candidate probe(s) meets selection criteria, its reverse complement is generated to enable binding to the positive-strand 16S rRNA. To complete the initiator probe design, an initiator sequence must then be tagged to the 3’ end of the reverse-complement probe, including to any other probes in the same set.

To design additional pan-bacterial 16S rRNA initiator probes (Eub841 and Eub1491), gene alignments were performed on a few bacteria from diverse phyla. Sites that were the most conserved were selected as candidate probes. Eub841 (paired with a second probe for expanded coverage) and Eub1491 were selected because they bind the same or greater number of bacterial taxa as Eub338 (>24729 sequences/26020 in our database; see **Data S2**). Together, these three probes target three different sites along the 16S rRNA (Eub338 between V2 and V3; Eub841 between V4 and V5; Eub1491 between V8 and V9) and cover 25954/26020 sequences in our database. The 66 bacteria not covered by the *Eubacteria* combined probe set are not found in the mammalian microbiota (**Data S2**). Cross-referencing all *Eubacteria* probes against human (Gene ID: 100008588) and mouse (Gene ID: 19791) 18S rRNA, as well as their mitochondrial 12S (human gene ID: 4549; mouse gene ID: 17724) or 16S rRNA (human gene ID: 4550; mouse gene ID: 17725), showed that the highest similarity was to the 12S mitochondrial rRNA with a minimum of three mismatches.

Sequences for all rRNA probes in this study are detailed in **Table S1**, and files displaying binding sites for each probe, including mismatches, are uploaded (**Data S2**). All probes within a probe set have the same initiator sequence and are pooled to maximize signal for each target bacteria. The non-split initiator design was used for probes targeting rRNA^38,39^ (**Figure S1**), and these sequences are detailed in **Table S2**.

For mRNA probes, a split initiator design was used (**Figure S1b**), which was created through the Özpolat Lab in situ HCR probe generator (rwnull/insitu_probe_generator on Github)^70^ for GFP, mCherry, and *Stm* ompA sequences. To compare ompA nucleotide identity between *Stm* (accession number FQ312003.1/locus tag SL1344_1010) and *E. coli* (accession number BA000007.3/locus tag ECs_1041), genes were aligned using CLC Main Workbench v8.1.

### Joint-Compositional Tensor Factorization (Joint-CTF)

Joint-CTF is a dimensionality reduction method for longitudinal multi-modal data based on tensor decomposition. It extends Compositional Tensor Factorization (CTF)^104^ and TEMPoral TEnsor Decomposition (TEMPTED)^126^, which are applied to a single data modality sampled over time. Joint-CTF simultaneously decomposes multiple longitudinal data modalities measured on the same set of subjects, while accommodating irregular sampling across subjects. For the analysis presented here, Joint-CTF was applied to two modalities: (1) absolute abundances of *Enterobacteriaceae* taxa and (2) antibody-bound taxa. Each data modality was log-transformed and z-scored to center the data. The normalized data were then reformatted into 3D tensors with axes corresponding to subjects, features, and time, and jointly decomposed. This decomposition generates (a) subject loadings, which embed each subject into a shared low-dimensional space of *n* principal components (PCs), capturing cross-modality variation; (b) feature loadings, which are modality-specific and quantify the relative contribution of individual features to each PC; and (c) temporal loadings, which are modality-specific and capture the dominant trajectories of features over time for each PC. To align time points across subjects with irregular sampling, original time points are projected into a high-dimensional embedding using kernel methods, creating a common temporal axis for all subjects.

### FlowJo analysis

Flow cytometry data were analyzed using FlowJo v10.10.0 (BD Biosciences). FlowJo UMAP v4.1.1 plugin was used, and consistent parameters were applied across all datasets (nearest neighbors = 5, minimum distance = 0.1, and components = 2). UMAPs were generated on concatenated patient files. FlowJo Cluster Explorer was used to visualize and present all UMAP data.

## QUANTIFICATION AND STATISTICAL ANALYSIS

### Absolute abundance calculations

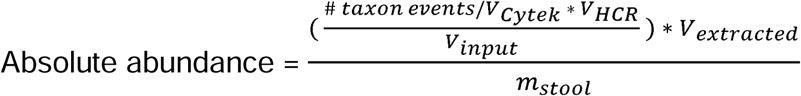

Where:

- # taxon events = number of events counted for taxon of interest
- V_Cytek_ = total volume (uL) of sample acquired by Cytek
- V_HCR_ = final volume (uL) of sample in 5X SSCT prior to running on flow cytometer
- V_input_ = volume (uL) taken from glycerol-stored microbiota used as input for testing
- V_extracted_ = volume (uL) of glycerol used for final resuspension of extracted microbiota
- m _stool_ = mass (mg) of stool from which microbiota was originally extracted

Example:

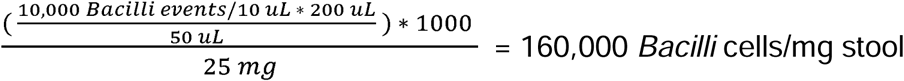

If there were 10,000 *Bacilli*+ events in our MicFLY-tested mouse stool sample, the Cytek volumetric function showed that 10 μL of sample was run through the cytometer, and 25 mg of a mouse stool pellet was originally processed, then according to the protocol, we would have 160,000 *Bacilli*+ cells/mg stool.

### Statistical analysis

All statistical analyses were performed in GraphPad Prism v10.4.1 unless otherwise indicated. Statistical tests used are in figure legends. Preterm infant group sizes were determined by the number of available samples within defined criteria in the cohort.

Stool testing and mice experiments were non-blinded. For comparisons between two groups, unpaired two-tailed Welch’s t-tests or Mann-Whitney U tests for nonparametric data were used. Paired two-tailed t-tests were applied to matched mouse stool samples. For comparisons among more than two groups, one-way ANOVA with Tukey’s multiple comparisons test was used. Two-way ANOVA with Tukey’s multiple comparisons test was used for mouse infection experiments comparing two time points with two taxa.

Microbiome Analyst^127^ generated PCoA plots on beta diversity with Bray-Curtis distance analysis; pairwise PERMANOVA with Benjamini-Hochberg false discovery rate (FDR) correction assessed significant differences in distances. Correlation analyses used Spearman correlation, with r and p-values reported. Lines in scatter plots represent the best-fit line from simple linear regression analysis. For multiple pairwise tests, nonparametric Mann-Whitney was adjusted with Holm–Sidak correction. Statistical significance is defined as p < 0.05.

---

**Supplemental Figure S1.**
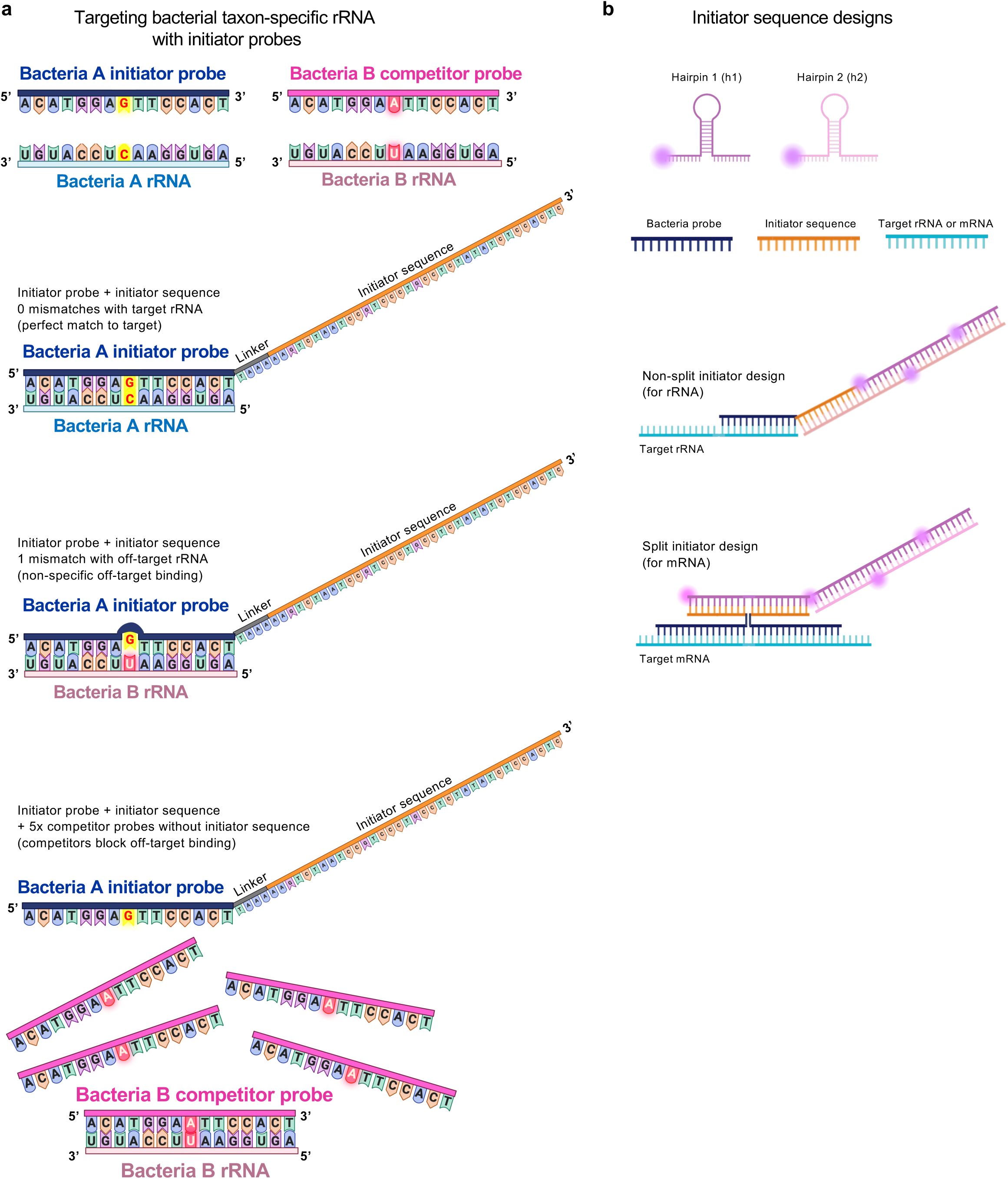
Initiator probe design a, Designing taxon-specific rRNA initiator probes with accompanying competitor probes to block highly similar but off-target rRNA. Competitor probes do not have an initiator sequence. b, Non-split initiator and split initiator sequence designs. Non-split initiator sequences are used for targeting rRNA, and split initiator sequences are used for targeting mRNA. Images made with BioRender.

**Supplemental Figure S2.**
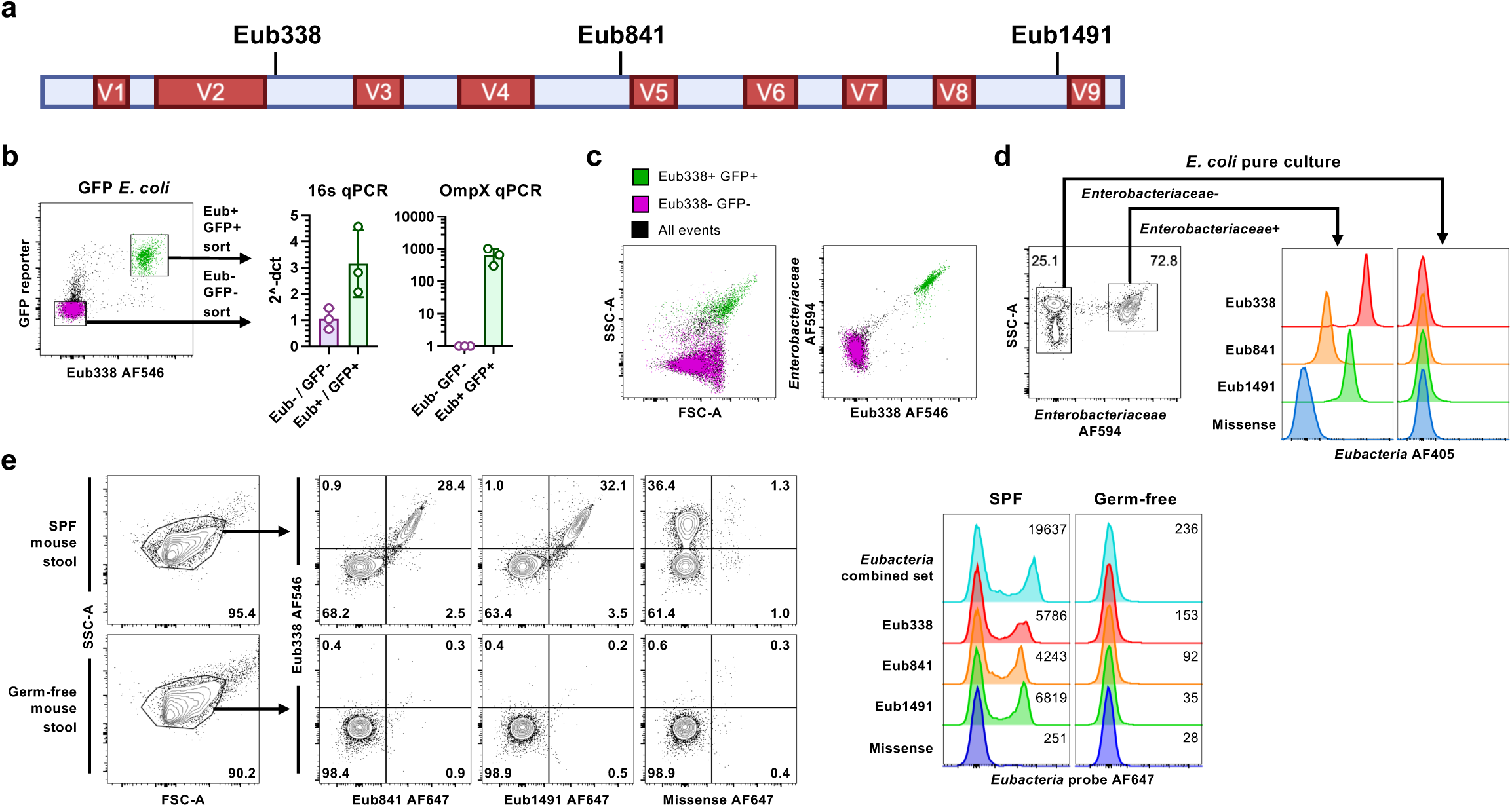
MicFLY effectively targets bacterial 16S rRNA a, Schematic of the 16S rRNA gene depicting the sites targeted by ‘pan-*Eubacteria*’ probes, made with BioRender. **b**, *E. coli* expressing arabinose-inducible Green Fluorescent Protein (GFP), induced by 0.01% arabinose, was analyzed with MicFLY. Flow plot (left) shows the gating strategy. GFP+ *Eubacteria*+ (green) and GFP*Eubacteria*-(purple) events were sorted into purified populations and assessed by quantitative PCR for the presence of 16s rRNA genes or *E. coli* ompX, relative to a buffer blank. Each dot represents an experimental replicate. c, Flow cytometry plots depicting the Forward and Side Scatter properties (left) or *Enterobacteriaceae* probe binding properties of GFP+ *Eubacteria*+ and GFP-*Eubacteria*-bacterial populations identified in b. d, *E. coli* was grown *in vitro* and used to test the three *Eubacteria* probes by MicFLY. The flow plot (left) shows the gating strategy to identify *E. coli* from debris and the histograms (right) depict the binding of the three *Eubacteria* probes to events from the two gates. e, MicFLY staining of fecal samples from specific pathogen free (SPF) or germ-free mice with the three *Eubacteria* probes. Flow plots (left) and histograms (right) demonstrate the staining of these probes within the two types of samples. Numbers in flow plots represent are percentage of events within that gate. Numbers in histograms represent mean fluorescence intensity. All histograms normalized to mode.

**Supplemental Figure S3.**
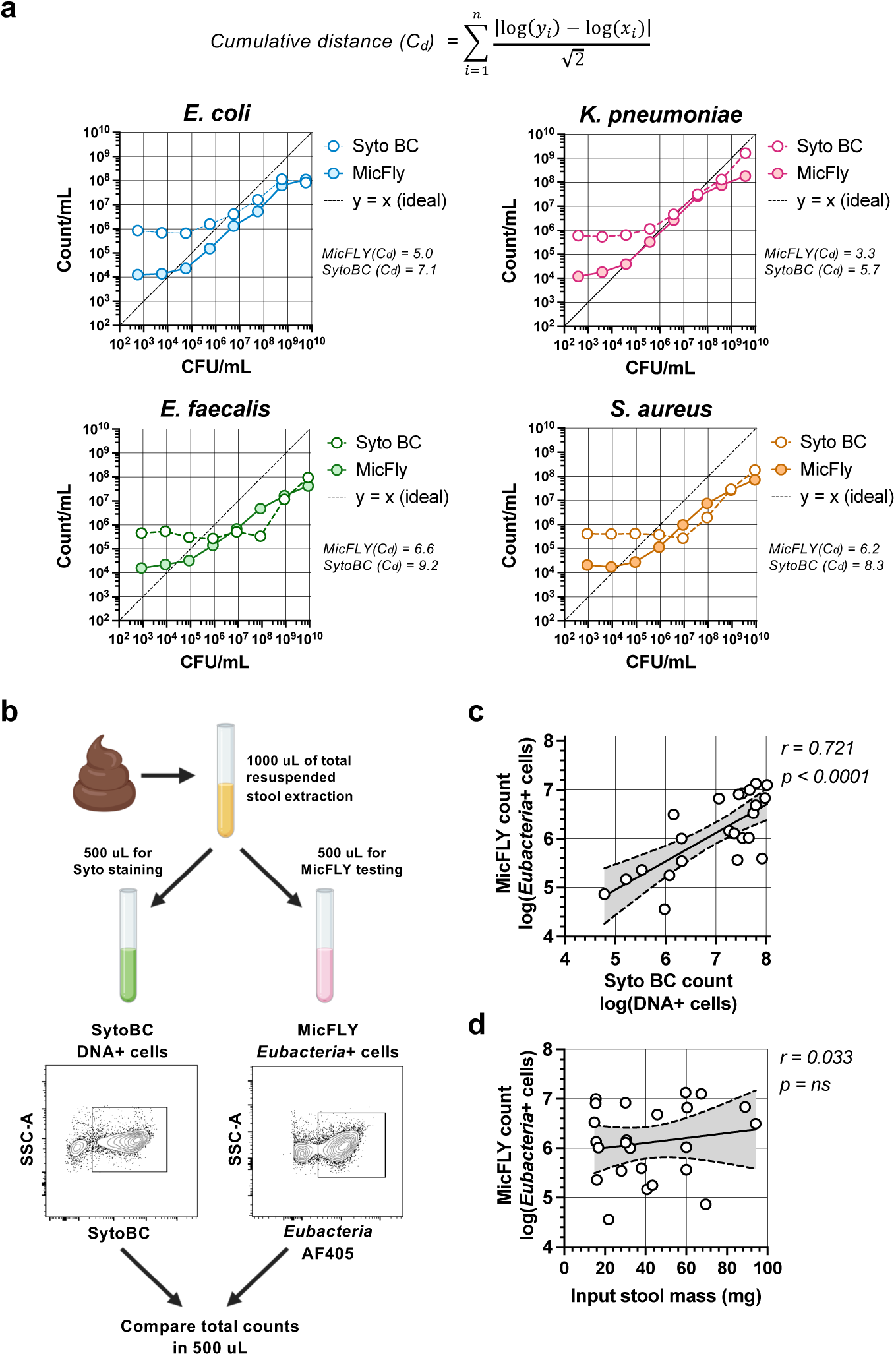
MicFLY permits bacterial quantitation. **a**, *In vitro* cultured bacterial samples were assessed for cellular abundance by dilution and plating or flow cytometry, performed with either MicFLY or the DNA stain, SytoBC. Graphs compare the cell number calculated by MicFLY (filled circles) and SytoBC (open circles) to growth on plates. The deviation of each technique from plating was calculated with the equation above and is provided to the right of each graph as cumulative distance (Cd) value. Dashed line represents where y values and x values are equal. b, Schematic for experiments comparing SytoBC and MicFLY enumeration of fecal samples. c, Correlation of bacterial abundances calculated from SytoBC and MicFLY data extracted from fecal samples from b. d, Correlation of bacterial abundances calculated from MicFLY to the mass of stool used for MicFLY testing. For (c,d), best-fit line from simple linear regression is displayed for log-transformed total abundance data, and shaded area represents 95% confidence interval; *r* and *p*-value from Spearman correlation

**Supplemental Figure S4.**
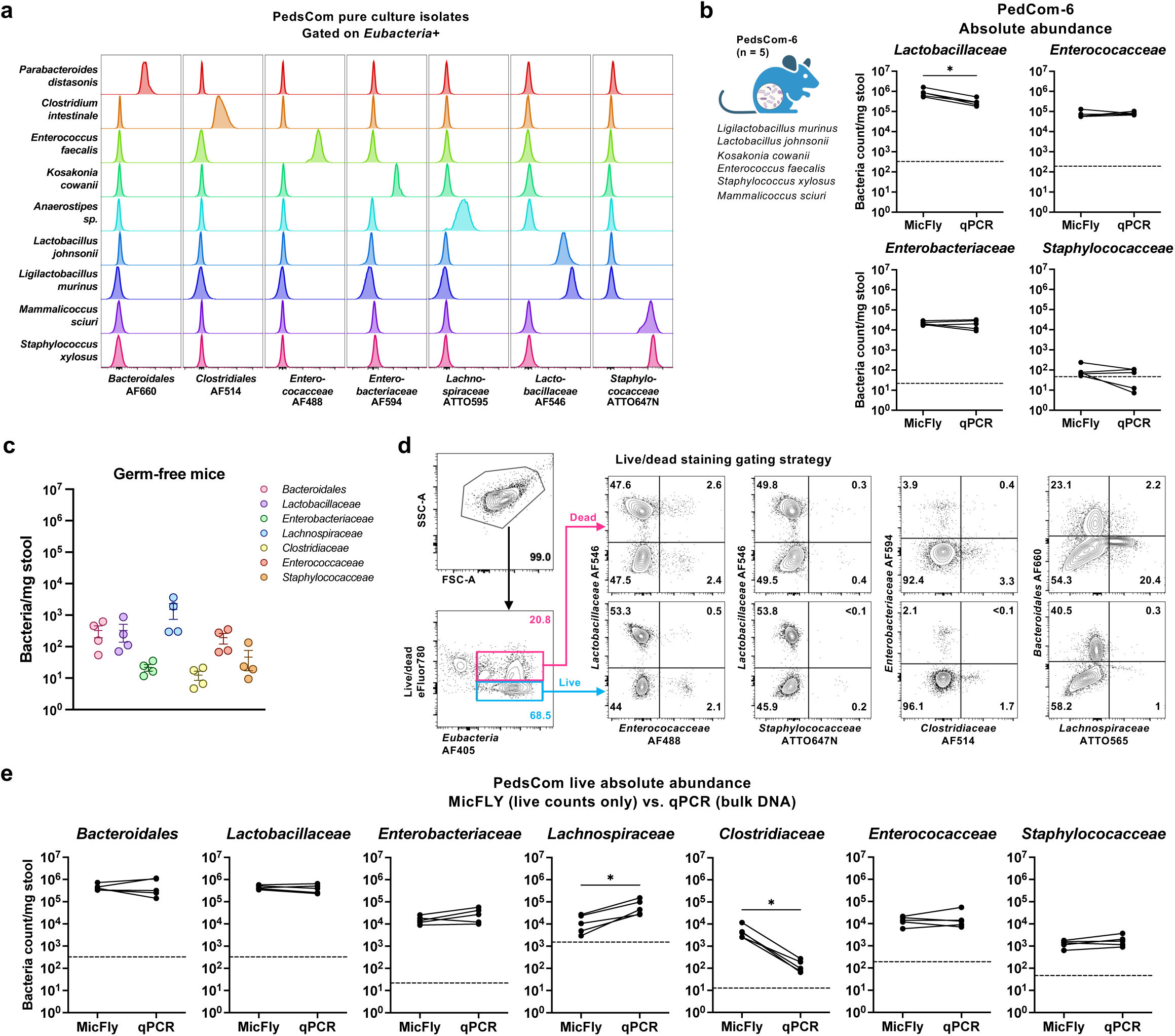
MicFLY can identify all bacteria in gnotobiotic mice. a, Multiplexed detection of 9 *in vitro* cultured bacterial species that together comprise ‘PedsCom’. Each column represents signal from a unique probe across different taxa. All probes were tested in the same panel and stained on individual taxa. Cells are gated on total bacteria (as in Fig. 1c). b, Absolute abundances of each bacterial family measured by MicFLY and qPCR from paired stool samples for 5 mice colonized with ‘PedsCom-6’ (described in image; left). Bacterial counts normalized to stool weight (mg), and statistics performed using a paired two-tailed t-test. Dashed line represents limit of detection, calculated as the mean absolute abundance in stool from germ-free mice (n = 4). p < 0.05 (*). **c**, The probes used for enumerating the taxa present in PedsCom were tested against fecal samples from four germ-free C57BL/6 mice. Number of positive events detected are graphed for each probe. Each dot represents a separate germ-free mouse. d, Flow cytometry plots depicting the gating strategy (left) and percent of each bacterial taxa within PedsCom-9 that is either live or dead, as determined by staining with fixable viability dye eFluor 780. e, Comparison of the absolute abundances of live bacteria (calculated by MicFLY) and all bacteria (calculated by qPCR) of family-level taxa from five mice colonized with the 9-member PedsCom. Bacterial counts were normalized to stool weight (mg), and statistical comparisons were performed using a paired two-tailed t-test. Dashed line represents limit of detection for that taxon, calculated as mean absolute abundance in stool from germ-free mice (*n = 4*). p < 0.05 (*).

**Supplemental Figure S5.**
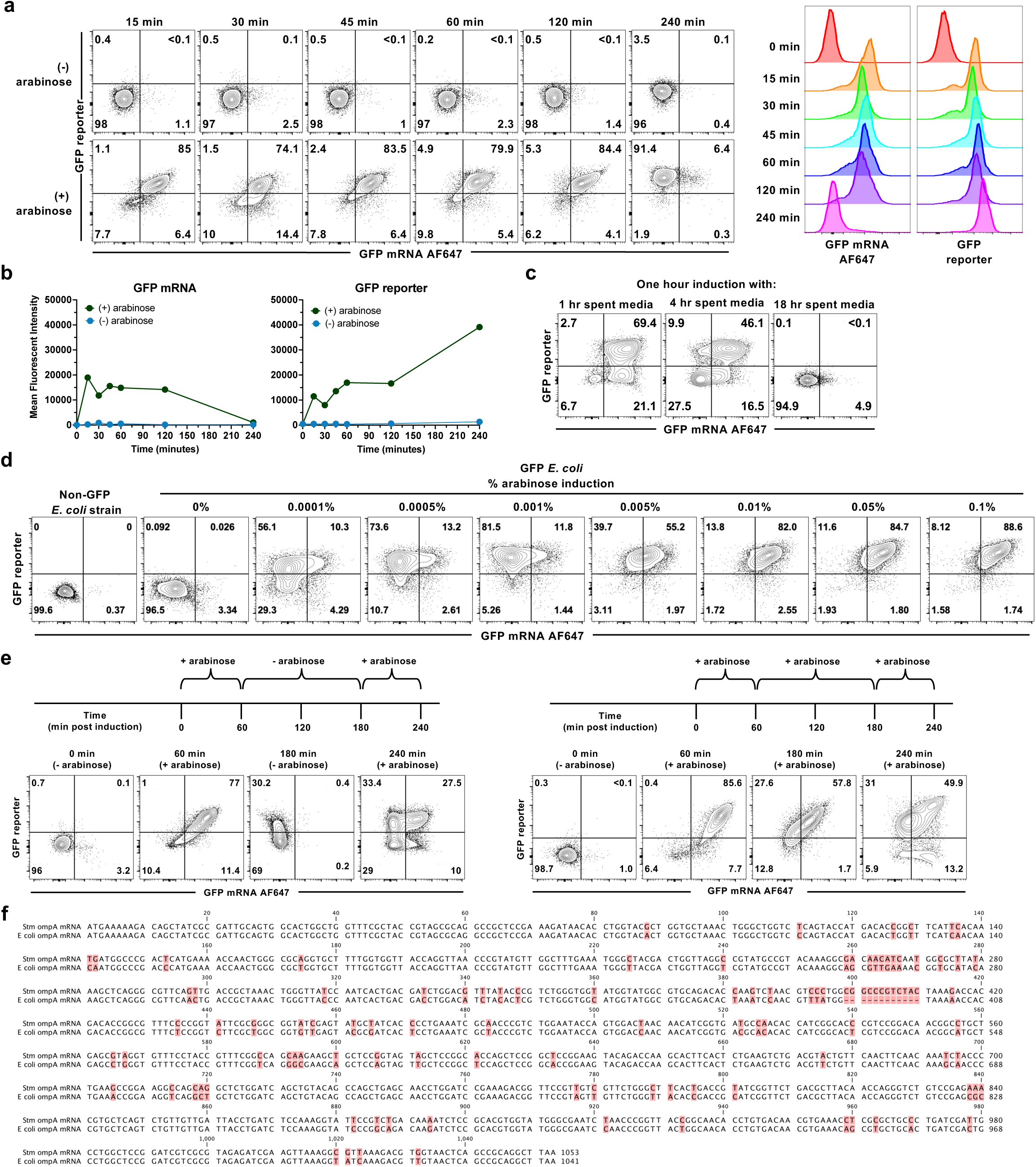
MicFLY can be used to identify mRNA expression levels in bacteria. a-e, A plasmid containing an arabinose-inducible Green Fluorescent Protein (GFP) molecule transformed into *E. coli*. a, Flow cytometry plots depicting GFP and GFP mRNA expression in *E. coli* induced with arabinose (0.01%; bottom row) or without arabinose as control (top row). Histograms (right) show the same data but allow comparison of relative expression over time. Bacteria are gated on *Eubacteria*/B17-AF405+ *Enterobacteriaceae*/B2-AF594+ cells. b, Graphs depict quantitation of a. c, Media was extracted from the 1-hour (60 minutes), 4-hour (240 minutes) and 18-hour (overnight) timepoints from (a) and used to activate GFP expression in new samples of *E. coli*. Shown is expression of GFP reporter protein and GFP mRNA (measured by MicFLY). d, Flow cytometric plots showing the response of GFP-expressing *E. coli* subpopulations to different concentrations of arabinose at 3 hours post-induction. Shown is expression of GFP reporter protein and GFP mRNA (measured by MicFLY). e, Flow cytometric plots depicting the effect of a time course of arabinose induction and removal on GFP-expressing *E. coli*. Shown is expression of GFP reporter protein and GFP mRNA (measured by MicFLY). **f**, Comparison of the sequence of the ompA gene from *Stm* and *E. coli*. Differences in the sequence of the genes is highlighted in red. Gene alignments made using CLC Main Workbench.

**Supplemental Figure S6.**
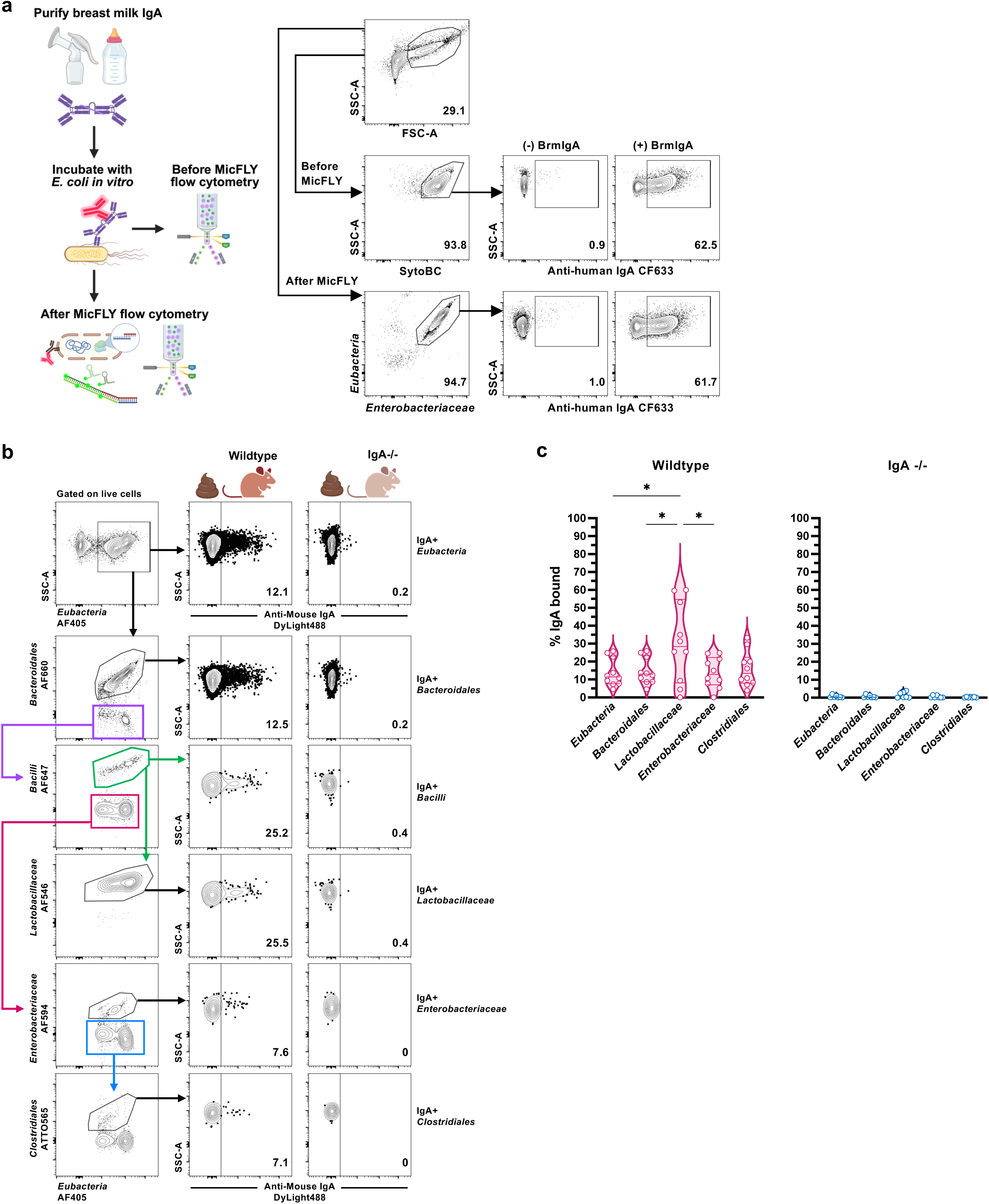
MicFLY retains surface antibody binding on bacteria. a, A schematic of the experiment is shown on the left (made with BioRender). *In vitro* cultured *E. coli* was incubated with IgA isolated from a breast milk sample, and then crosslinked/fixed with anti-fluorescent IgA antibody (step 1b-2 in Fig. 1a). Bacteria incubated without breast milk IgA, but with secondary fluorescent antibody, was used as a control. IgA binding was quantified before and after MicFLY to assess any loss in binding due to the protocol (step 3 onward in Fig. 1a). Shown is the gating strategy (flow cytometry plots on the left) and level of IgA binding to the bacteria (flow cytometry plots on the right). b, Fecal samples were isolated from C57BL/6 and IgA-/- mice and analyzed by MicFLY. Flow cytometric plots on the left show the gating strategy. Flow cytometric plots on the right show percentage of IgA binding to various taxa within the mouse microbiota. Center column = C57BL/6; Right column = IgA-/- **c**, Quantification of b. Statistical tests were performed using ordinary one-way ANOVA with Tukey’s multiple comparisons test. p < 0.05 (*).

**Supplemental Figure S7.**
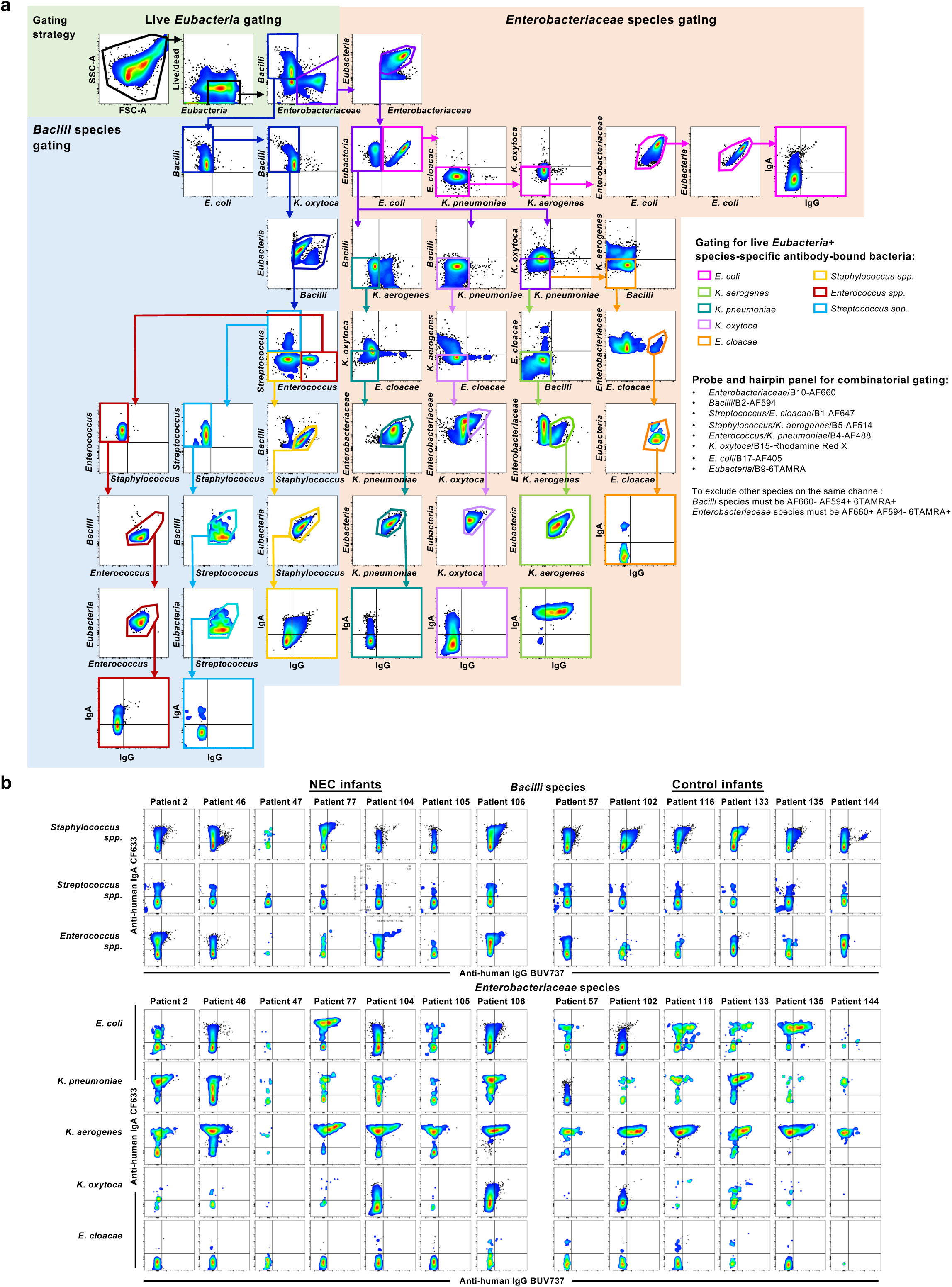
Gating strategy for combinatorial spectral flow cytometry to identify live antibody-bound bacterial taxa in infant stool. a, Gating strategy used to identify live bacterial species in stool samples from control or NEC infants using MicFLY. Flow cytometry plots are representative examples from different patient-specific concatenated files, as no single infant contained all taxa shown. Negative gating for all other taxa was applied to each genus/species to ensure it is positive only for its own probe within the *Enterobacteriaceae* or *Bacilli* parent gate. We stacked fluorophores by using combinatorial gating. For example, each *Bacilli* species is negative for *Enterobacteriaceae*/B10-AF660, but positive for *Bacilli*/B2-AF594, and each *Enterobacteriaceae* species is positive for *Enterobacteriaceae*/B10-AF660 but negative for *Bacilli*/B2-AF594. Thus, species-specific probes can be doubled up, as long as they are in different families/orders (e.g., *Staphylococcus/K*. *aerogenes*/B5-AF514; *Enterococcus/K. pneumoniae*/B4-AF488). Colored boxes highlight strategy used to gate for each taxon. b, Species-specific IgA and IgG binding patterns across NEC and control infants. Flow cytometry plots were derived from concatenated files of all samples for each individual patient. Shown is IgA/IgG-bound taxa for every patient, based on anti-human IgA (CF633, y-axis) and IgG (BUV737, x-axis). Pseudocolor smoothing with large dot size was applied to improve visualization of low-abundance taxa.

**Supplemental Figure S8.**
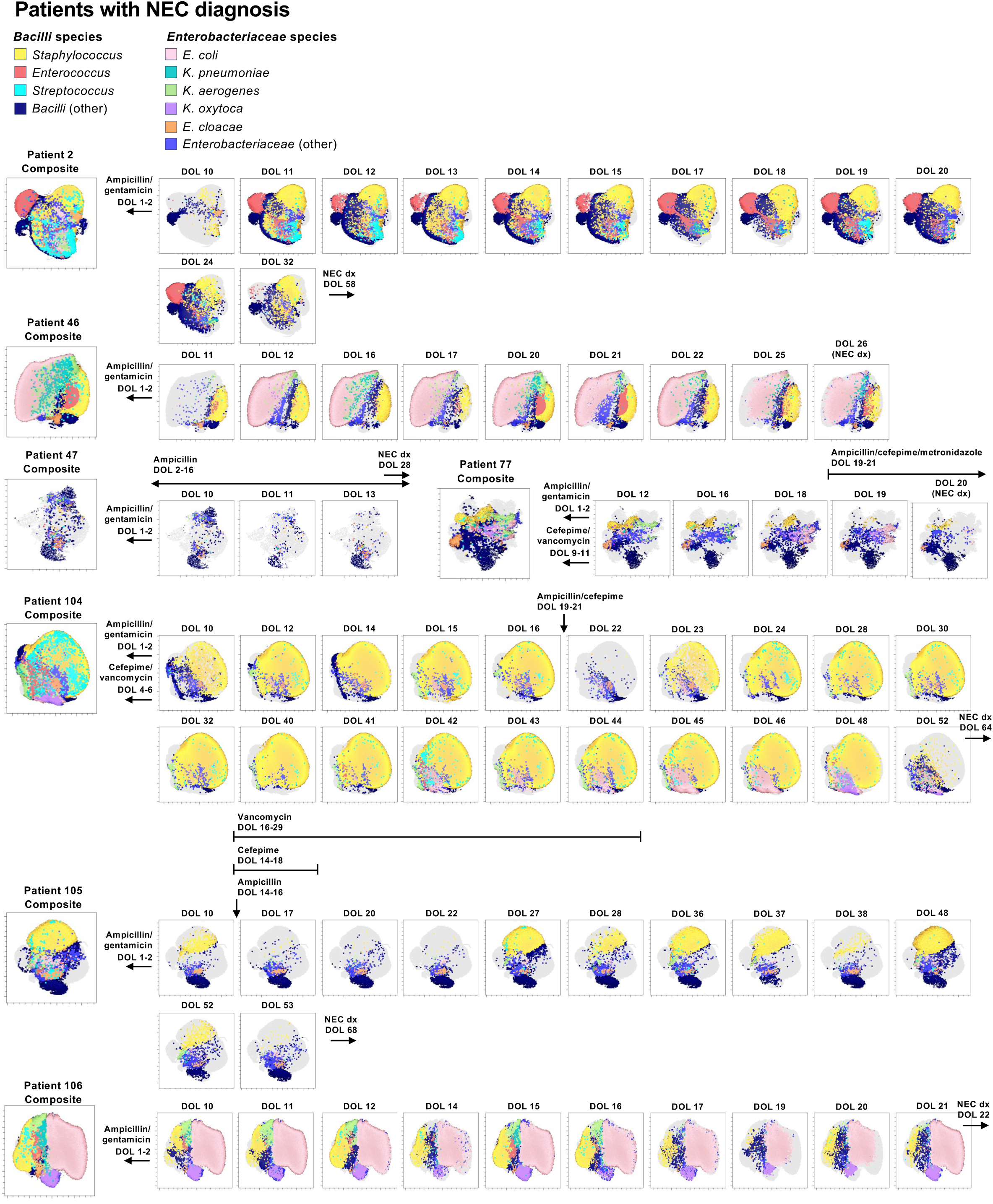
Day-to-day UMAP plots of NEC-diagnosed infants. UMAP plots of composition for both *Bacilli* and *Enterobacteriaceae* species for all stool samples from individual preterm infants who develop necrotizing enterocolitis (NEC). Composite UMAPs (left) represent concatenated data from all available samples per patient, overlaid on live *Eubacteria*/B9-TAMRA+ gated populations. Clusters are color-coded according to taxon, with gray area on the composite UMAP representing unclassified bacteria. UMAPs for each DOL was overlaid on the patient composite UMAP, with gray areas on sample-specific UMAPs corresponding to bacteria not covered on composite. Plots generated using FlowJo’s UMAP plugin (5 nearest neighbors, minimum distance 0.1, 2 components) and visualized with Cluster Explorer. Each dot represents a single bacterial cell, color-coded according to species. Antibiotic administration is indicated above each time point where applicable. DOL at which NEC will be diagnosed is marked with a black arrow labeled “NEC dx.”

**Supplementary Figure 9.**
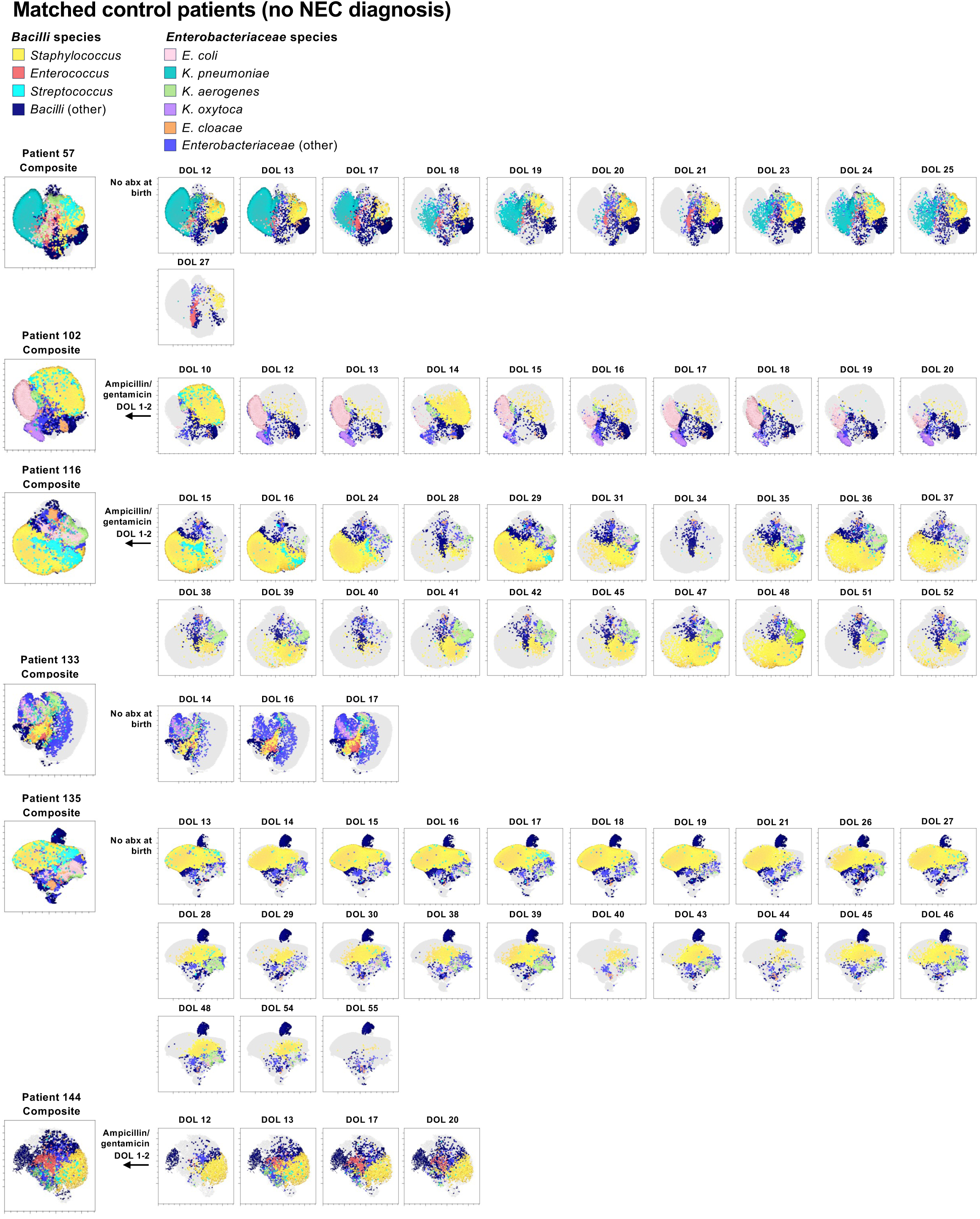
Day-to-day UMAP plots of matched control infants. UMAP plots are displayed for all stool samples from individual matched control preterm infants. Parameters/visualization exactly as in Supplementary Figure 8.

**Supplemental Figure S10.**
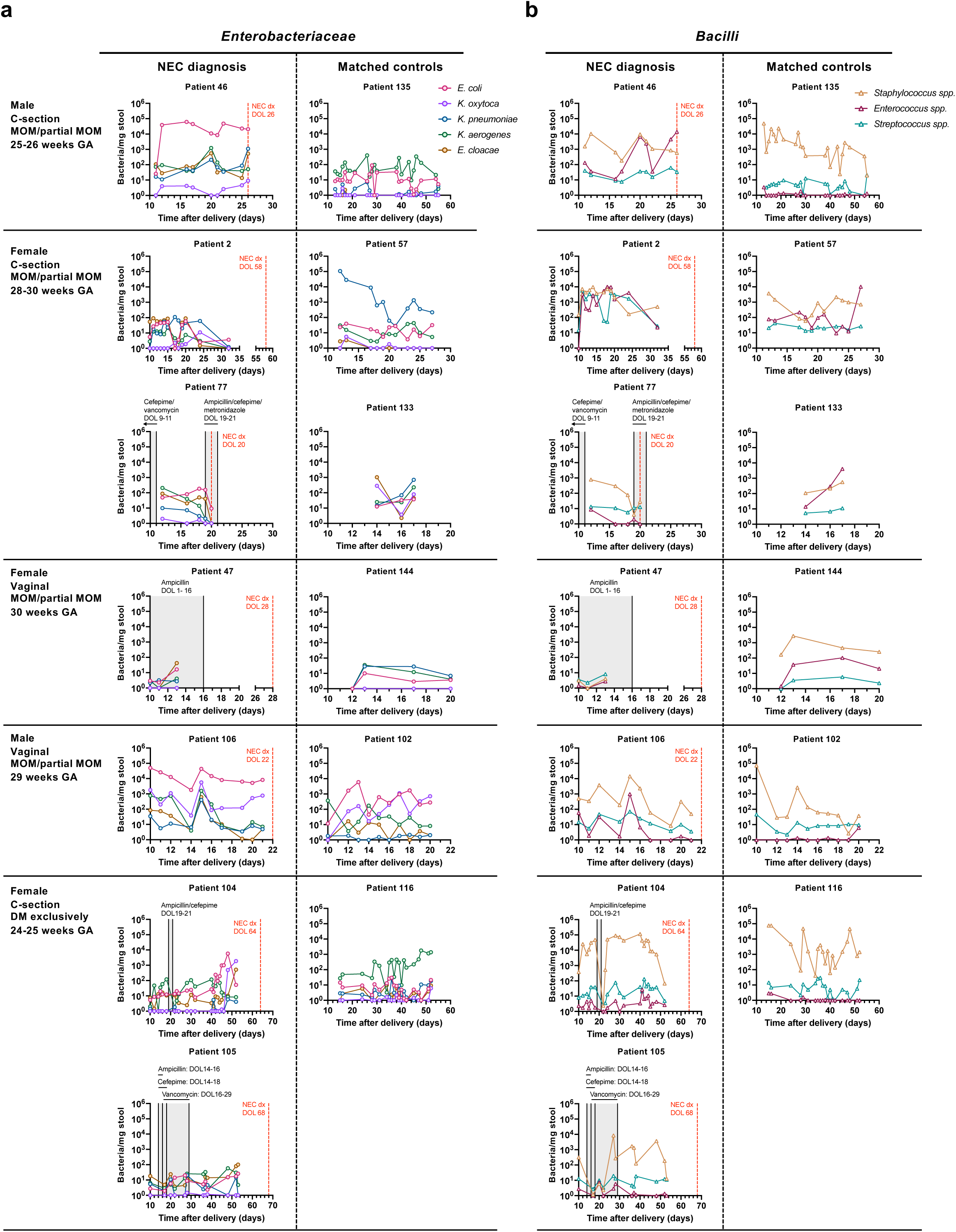
Longitudinal absolute abundance of *Enterobacteriaceae* and *Bacilli* species in NEC and control groups. a, Total counts for *Enterobacteriaceae* species; b, total counts for *Bacilli* species. Plots show absolute abundance, measured by MicFLY of each species in stool samples from preterm infants who developed NEC (left) versus matched controls (right). Each panel displays data from a single infant over time postdelivery (x-axis), with DOL of future NEC diagnosis marked by a red dashed line. Antibiotic exposure windows are shaded in gray. Different line colors represent each species. Patients were matched by gestational age (within 2 weeks), sex, delivery mode, and feeding type (mother’s own milk [MOM] vs. donor milk [DM]). NEC patients (n=7) and matched controls (n=6) were selected from the prospective longitudinal cohort shown in Figure 5.

**Supplemental Figure S11.**
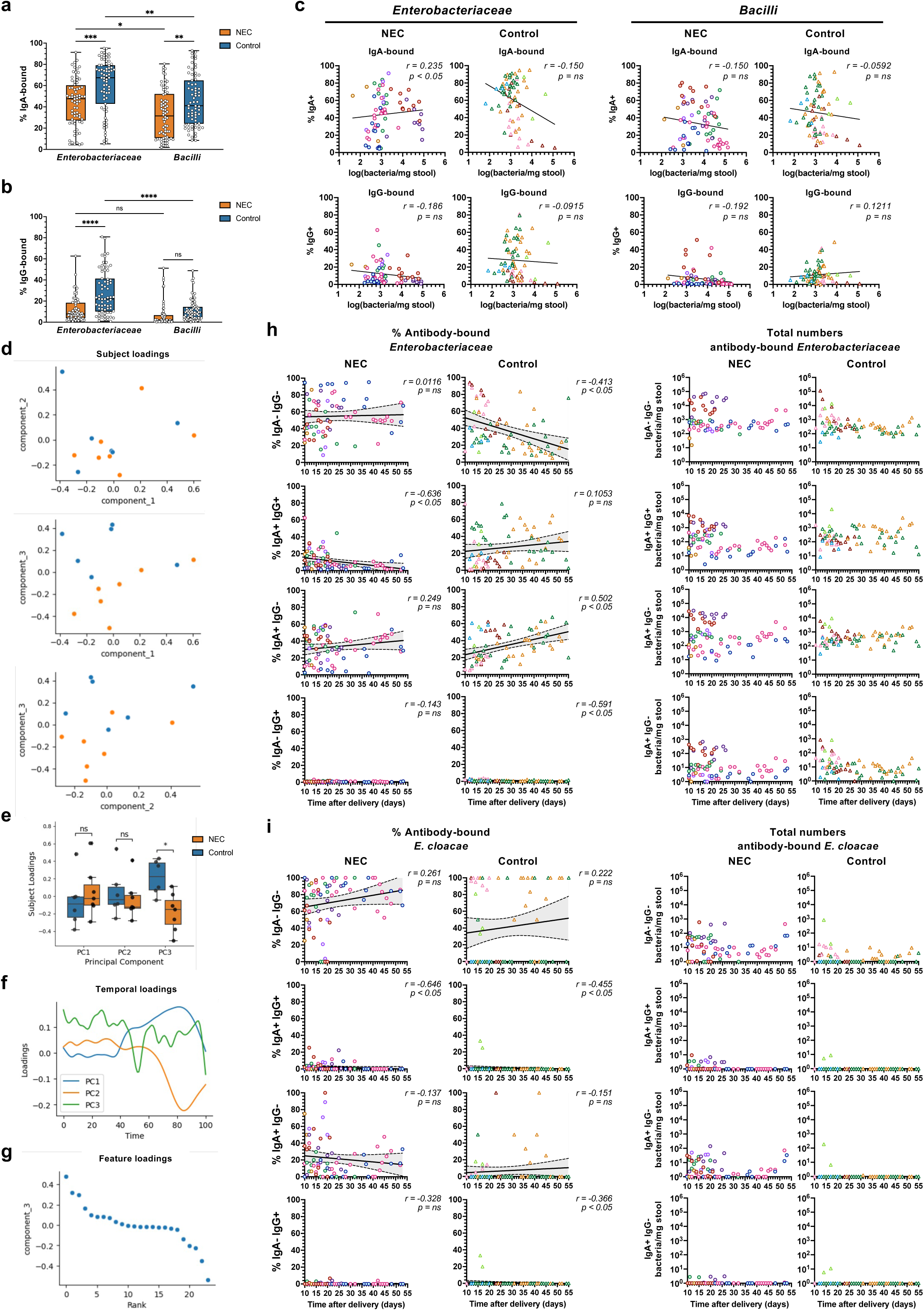
Correlation of antibody binding to infant intestinal bacteria to the development of NEC. a, Graph depicting the percent of IgA-bound *Enterobacteriaceae* and *Bacilli* species in stool samples from NEC (n=7) and control (n=6) infants, as measured by MicFLY. Box plots show median, interquartile range, and minimum to maximum values. Statistical tests were performed using ordinary one-way ANOVA with Tukey’s multiple comparisons test. p < 0.05 (*), p < 0.01 (**), p < 0.001 (***), p < 0.0001 (****). b, Percent of IgG-bound *Enterobacteriaceae* and *Bacilli* species as in panel a. c, Graphs of the correlation between bacterial abundance (log-transformed) and percent IgA or IgG bound bacteria within each taxon, measured by MicFLY, for *Enterobacteriaceae* and *Bacilli* in NEC and control groups. Best-fit line from simple linear regression is displayed; *r* and *p*-value from Spearman correlation. Each dot represents a single sample, and each color corresponds to an individual patient. Repeated colors indicate multiple samples per patient. d-g, Joint-CTF multi-modal dimensionality reduction using three principal components (PCs) derived from absolute abundances of Enterobacteriaceae species and antibody binding over time. d, Subject loadings, where each point represents an infant’s overall multi-modal profile embedded into a shared low-dimensional space. Infants from the NEC (orange) and control (blue) groups are shown across pairwise combinations of the three PCs. **e**, Separation of NEC and control groups along each PC from panel **d**. Group differences were assessed using the Mann-Whitney U test; p < 0.05 (*). f, Temporal loadings capturing dominant feature trajectories over time. The x-axis represents a highdimensional projection of DOL, where original DOL values (1-55) were projected and rescaled to a common 1-100 scale, enabling alignment of irregularly sampled infants. g, Feature loadings on PC3, reflecting the relative contributions of features to group separation. Based on panel e, negative PC3 values (y-axis) associate with NEC, while positive PC3 values with controls. h, Graph comparing DOL and percent (left) or absolute abundance (right) of antibody-bound *Enterobacteriaceae* in NEC and control patients. Antibody binding is split between IgG- and IgA-bound bacteria, as indicated. Best-fit line from simple linear regression is displayed, and shaded area represents 95% confidence interval; *r* and *p*-value from Spearman correlation. Same samples and color coding as in panel c. i, Longitudinal display of DOL and percent (left) or absolute abundance (right) of antibody-bound *E. cloacae* in NEC and control patients, exactly as in panel h.

