## Supplemental Notes and Tables for "Single-cell quantification of the microbiota by flow cytometry: MicFLY"

**Supplemental Note S1: MicFLY-tested fluorophores**

****All fluorophores are spectrally compatible unless otherwise indicated*

Hairpin fluorophores that work with MicFLY

- Alexa Fluor 660
- Alexa Fluor 647
- Alexa Fluor 594
- Alexa Fluor 546 (cannot be used with 6-TAMRA on spectral panel)
- Alexa Fluor 488 (cannot be used with DyLight 488 on spectral panel)
- Alexa Fluor 514
- Alexa Fluor 405
- Rhodamine Red X (cannot be used with ATTO565 on spectral panel)
- 6-TAMRA (cannot be used with Alexa Fluor 546 on spectral panel)
- Texas Red (cannot be used with Alexa Fluor 594 on spectral panel)
- ATTO490LS
- ATTO565 (cannot be used with Rhodamine Red X on spectral panel)
- ATTO647N

Hairpin fluorophores that have too low signal and **do not** work well with MicFLY

- Alexa Fluor 430
- ATTO532
- ATTO425

Antibody fluorophores that work with MicFLY

- DyLight488 (cannot be used with Alexa Fluor 488 on spectral panel)
- CF633
- BUV737
- VioGreen (has not been tested with BUV737 on same spectral panel)

Antibody fluorophores that have been tested and **do not** survive MicFLY protocol

- Protein-based fluorophores (e.g., APC, PE)
- Tandem conjugates (e.g., PE-Cy7, PE-APC, PE-Vio770, etc.)

Fixable viability dye for live/dead staining that works with MicFLY

- eFluor780

### Supplemental Table S1: rRNA oligonucleotide probe sequences used in this study

| Target bacteria | | Associated initiator sequences used in this study (see Supplementary Table 2) | Probe name | Target taxonomic level | rRNA (16s or 23s) | Probe sequence (5'-3') | Reference |
|---|---|---|---|---|---|---|---|
| *Eubacteria* probe set | | B1 – B17 | Eub338-I | Kingdom | 16s | GCTGCCTCCCGTAGGAGT | PMID: 2200342 |
| | | | Eub338-II | Kingdom | 16s | GCAGCCACCCGTAGGTGT | PMID: 10553296 |
| | | | Eub338-III | Kingdom | 16s | GCTGCCACCCGTAGGTGT | PMID: 10553296 |
| | | | Eub841-I | Kingdom | 16s | CTACCAGGGTATCTAATCC | This study |
| | | | Eub841-II | Kingdom | 16s | CTACCGGGGTATCTAATCC | This study |
| | | | Eub1470 | Kingdom | 16s | GACGGGCGGTGTGTACAAG | This study |
| *Enterobacteriaceae* probe set | | B2, B10 | EnterobactD | Family | 16s | TGCTCTCGCGAGGTCGCTTCTCTT | PMID: 12067375 |
| | | | Ent190-I | Family | 16s | CCCCCTCTTTGGTCTTGC | This study |
| | | | Ent190-II | Family | 16s | CCCCCACTTTGGTCTTGC | This study |
| | | | Ent190-III | Family | 16s | GAGCCCCTGCTTTGGTCCGTA | This study |
| | | | Enterobact1317-I | Family | 16s | GACGCACTTTATGAGGTC | This study |
| | | | Enterobact1317-II | Family | 16s | GACATACTTTATGAGGTC | This study |
| | | | Enterobact1317-III | Family | 16s | GACAGACTTTATGAGTTC | This study |
| *Bacilli* probe set | | B2, B3 | Firmicutes-A | Class | 16s | TGGAAGATTCCCTACTGC | PMID: 10390869 |
| | | | Firmicutes-B | Class | 16s | CGGAAGATTCCCTACTGC | PMID: 10390869 |
| | | | Firmicutes-C | Class | 16s | CCGAAGATTCCCTACTGC | PMID: 10390869 |
| *Lactobacillaceae* probe set | | B3 | Lacto663 | Family | 16s | ACATGGAGTTCCACT | This study |
| | | | Lacto1261 | Family | 16s | GACTCGTTGTACCGTCCATT | This study |
| | | No initiator sequence | Lact663 competitor | | 16s | ACATGGAATTCCACT | This study |
| *Staphylococaceae* probe set | | B1 | Staph1261-I | Family | 16s | TGCCCTTTGTATTGTCCATT | This study |
| | | | Staph1261-II | Family | 16s | TACCCTTTGTATTGTCCATT | This study |
| *Mammalicoccus sciuri* (used in PedsCom mice in *Staphylococacceae* set) | | B1 | Msciuri1261 | Species | 16s | TGCCCTTTGTATTATCCATT | This study |
| *Enterococaceae* probe set | | B2, B4 | EC1261-I | Family | 16s | GACTCGTTGTACTTCCCATT | This study |
| | | | EC1261-II | Family | 16s | AACTCGTTGTACTTCCCATT | This study |
| | | No initiator sequence | EC1261-I competitor | | 16s | GACTCGTTGTACTTGCCATT | This study |
| | | | EC1261-II competitor | | 16s | GACTCGTTGTACTGCCCATT | This study |
| *Bacteroidales* probe set | | B10, B11 | Bact982 | Order | 16s | ATGTTCCTCCGCTTGTGC | This study |
| | | | CFB1082 | Order | 16s | TGGCACTTAAGCCGACAC | PMID: 10879984 |
| *Clostridiales* probe set | *Lachnospiraceae* probe set | B15 | Erec482-I | Family | 16s | GCTTCTTAGTCAGGTACCG | PMID: 9726880 |
| | | | Erec482-II | Family | 16s | GCTTTTTAGTCAGGTACCG | This study |
| | | | Lachno501-I | Family | 16s | ATAGAAGTTTACATACCGA | This study |
| | | | Lachno501-II | Family | 16s | ATAGAGCTTTACATACCGA | This study |
| | *Ruminococcaceae* probe set | B15 | Clep866-I | Family | 16s | GGTGGATTACTTATTGTG | PMID: 15946290 |
| | | | Clep866-II | Family | 16s | GGTGGGATACTTATTGTG | This study |
| | *Clostridiaceae* probe set | B5, B15 | Chis150-I | Family | 16s | TTATGCGGTATTAATCTTCCTTT | PMID: 9726880 |
| | | | Chis150-II | Family | 16s | TTATGCGGTATTAATCTCCCTTT | This study |
| | | No initiator sequence | Chis150-I competitor-I | | 16s | TTATGGGGTATTAATCTTCCTTT | This study |
| | | | Chis150-I competitor-II | | 16s | TTATGAGGTATTAATCTTCCTTT | This study |
| | | | Chist150-II competitor | | 16s | TTATGGGGTATTAATCTCCCTTT | This study |
| | *Peptostreptococaceae* probe set | B1, B15 | Clt135-I | Family | 16s | GTTATCCGTGTGTACAGGG | PMID: 9726880 |
| | | | Clt135-II | Family | 16s | GTTATCCATGTGTATAGGG | This study |
| | *Finegoldia magna* | B15 | PTPHC454 | Species | 16s | GGACAGAACTTTACGATACG | This study |
| *Negativicutes* probe set | | B5, B9 | Prop853 | Class | 16s | ATTGCGTTAACTCCGGCAC | PMID: 16000778 |
| | | | Neg1243 | Class | 16s | CCGTGCACCACCTGTTTTC | This study |
| | | | Neg1243 competitor | Class | 16s | CCGTGCAGCACCTGTTTTC | This study |
| *E. coli* probe set | | B3, B17 | Ecoli 16s-460 | Species | 16s | GAGCAAAGGTATTAACTTTACTC | This study |
| | | | Ecoli 23s-342 | Species | 23s | ATGTGCATTTTTGTGTACGGGGC | This study |
| | | No initiator sequence | Ecoli 23s competitor-I | | 23s | CAGACTCTGGGCTGCTCC | This study |
| | | | Ecoli 23s competitor-II | | 23s | ATGTGCATTTTCGTGTACGGGGC | This study |
| *Salmonella typhimurium* | | B4 | Sal1 | Species | 23s | ACAGCACATGCGCTTTTGTG | PMID: 12747677 |
| | | | Sal3 | Species | 23s | AATCACTTCACCTACGTG | PMID: 12747677 |
| *Gammaproteobacteria* | | B2, B5 | Gam42A | Class | 23s | GCCTTCCCACATCGTTT | PMID: 9212435 |
| | | No initiator sequence | Gam42A competitor | | 23s | GCCTTCCCACTTCGTTT | PMID: 9212435 |
| *Actinobacteria* | | B10 | HGC69A | Phylum | 23s | TATAGTTACCACCGCCGT | PMID: 8000548 |
| *Klebsiella pneumoniae* probe set | | B4 | Kpn 23s-1700 | Species | 23s | CCTACACACCAGCGTGCC | PMID: 10655393 |
| | | | Kpn 16s-464 | Species | 16s | CGATGAGGTTATTAACCTCATCG | This study |
| *Klebsiella oxytoca* probe set | | B5, B15 | Koxy 23s-1700 | Species | 23s | CACTTACCATCAGCGTGCC | This study |
| | | | Koxy 23s-342 | Species | 23s | CTGTGTGTTTTAGTGTACGGGAG | This study |
| | | | Koxy 16s-464 | Species | 16s | GAATAAGGTTATTAACCTCACTC | This study |
| | | | Koxy 23s-1700 competitor | | 23s | CACCTACCATCAGCGTGCC | This study |
| *Klebsiella aerogenes* | | B2 | Kaero 16s-464 | Species | 16s | CGCCAAGGTTATTAACCTTAACG | This study |
| | | No initiator sequence | Kaero 16s-464 competitor | | 16s | CGCTAAGGTTATTAACCTTAACG | This study |
| *Enterobacter cloacae* | | B1 | Ecloacae 23s-342 | Species | 16s | CTGTGTGTTTTCGTGTACGGGAC | This study |
| | | | Ecloacae 23s-342 competitor | | 16s | CTGTGCGTTTTCGTGTACGGGAC | This study |
| *Streptococcus spp.* | | B1 | Str-Strep | Genus | 16s | CACTCTCCCCTTCTGCAC | PMID: 10917153 |
| | | No initiator sequence | Str competitor-I | | | CACTCTCCTCTTCTGCAC | This study |
| | | | Str competitor-II | | | CACTCTCCCCTCCTGCAC | This study |
| | | | Str competitor-III | | | CACTATCCCCTTCTGCAC | This study |
| *Fusobacterium* | | B4 | FUSO | Genus | 16s | CTAATGGGACGCAAAGCTCTC | PMID: 12724371 |
| *Akkermansia muciniphila* | | B1 | MUC-1437 | Species | 16s | CCTTGCGGTTGGCTTCAGAT | PMID: 18083887 |

#### Supplemental Table S2: Initiator and hairpin sequences

| Initiator sequence name | Initiator sequence (bold) + linker sequence (lowercase)<br><br>(5'-3'; non-split design; tagged on 3'end of initiator rRNA probe; see Supplementary Table 1) | Associated hairpin pair | Hairpin sequence (5'-3') | Hairpin fluorophore used in this study (conjugated to 5'end of h1 and h2) | Reference |
|---|---|---|---|---|---|
| B1 | aaaaa **GCATTCTTTCTTGAGGAGGGCAGCAAACGGGAAGAG** | B1h1 | GAGGAGGGCAGCAAACGGGAAGAGTCTTCCTTTACGCTCTT<br>CCCGTTTGCTGCCCTCCTCAAGAAAGAATGC | Alexa Fluor 647 | PMID: 24712299 |
| | | B1h2 | CGTAAAGGAAGACTCTTCCCGTTTGCTGCCCTCCTCGCATTC<br>TTTCTTGAGGAGGGCAGCAAACGGGAAGAG | | |
| B2 | aaaaa **AGCTCAGTCCATCCTCGTAAATCCTCATCAATCATC** | B2h1 | CCTCGTAAATCCTCATCAATCATCCAGTAAACCGCCGATGAT<br>TGATGAGGATTTACGAGGATGGACTGAGCT | Alexa Fluor 594 | PMID: 24712299 |
| | | B2h2 | GGCGGTTTACTGGATGATTGATGAGGATTTACGAGGAGCTC<br>AGTCCATCCTCGTAAATCCTCATCAATCATC | | |
| B3 | taaa **AAAGTCTAATCCGTCCCTGCCTCTATATCTCCACTC** | B3h1 | GTCCCTGCCTCTATATCTCCACTCAACTTTAACCCGGAGTGG<br>AGATATAGAGGCAGGGACGGATTAGACTTT | Alexa Fluor 546 | PMID: 24712299 |
| | | B3h2 | CGGGTTAAAGTTGAGTGGAGATATAGAGGCAGGGACAAAGT<br>CTAATCCGTCCCTGCCTCTATATCTCCACTC | | |
| B4 | atttt **CACATTTACAGACCTCAACCTACCTCCAACTCTCAC** | B4h1 | CCTCAACCTACCTCCAACTCTCACCATATTCGCTTCGTGAGA<br>GTTGGAGGTAGGTTGAGGTCTGTAAATGTG | Alexa Fluor 488 | PMID: 24712299 |
| | | B4h2 | GAAGCGAATATGGTGAGAGTTGGAGGTAGGTTGAGGCACAT<br>TTACAGACCTCAACCTACCTCCAACTCTCAC | | |
| B5 | atttt **CACTTCATATCACTCACTCCCAATCTCTATCTACCC** | B5h1 | ATTGGATTTGTAGGGTAGATAGAGATTGGGAGTGAGCACTTC<br>ATATCACTCACTCCCAATCTCTATCTACCC | Alexa Fluor 514<br>or<br>Alexa Fluor 405 | PMID: 24712299 |
| | | B5h2 | CTCACTCCCAATCTCTATCTACCCTACAAATCCAATGGGTAG<br>ATAGAGATTGGGAGTGAGTGATATGAAGTG | | |
| B9 | atttt **CACGTATCTACTCCACTCTCAGCACACTCCCAACCC** | B9h1 | CCACTCTCAGCACACTCCCAACCCTACTACAAGCTCGGGTT<br>GGGAGTGTGCTGAGAGTGGAGTAGATACGTG | ATTO490LS<br>or<br>6-TAMRA | PMID: 38966983 |
| | | B9h2 | GAGCTTGTAGTAGGGTTGGGAGTGTGCTGAGAGTGGCACGT<br>ATCTACTCCACTCTCAGCACACTCCCAACCC | | |
| B10 | atatt **CCTCAAGATACTCCTCTACCTACTCGACTACCCTAG** | B10h1 | CCTCTACCTACTCGACTACCCTAGCCGTAACTTCACCTAGGG<br>TAGTCGAGTAGGTAGAGGAGTATCTTGAGG | Alexa Fluor 660 | PMID: 38966983 |
| | | B10h2 | GTGAAGTTACGGCTAGGGTAGTCGAGTAGGTAGAGGCCTCA<br>AGATACTCCTCTACCTACTCGACTACCCTAG | | |
| B11 | aaaaa **CGCTTAGATATCACTCCTACGTCGACCACACTCATC** | B11h1 | ACTCCTACGTCGACCACACTCATCCTGCATGTTCCCGATGA<br>GTGTGGTCGACGTAG GAGTGATATCTAAGCG | ATTO647N | PMID: 38966983 |
| | | B11h2 | GGGAACATGCAGGATGAGTGTGGTCGACGTAGGAGTCGCTT<br>AGATATCACTCCTA CGTCGACCACACTCATC | | |
| B15 | atttt **CAGATTAACACACCACAAGGTATCTCGAACACTCTC** | B15h1 | CCACAAGGTATCTCGAACACTCTCCAAATTGGCTACGAGAG<br>TGTTCGAGATACCTTGTGGTGTGTTAATCTG | Rhodamine Red X<br>or<br>ATTO565 | PMID: 38966983 |
| | | B15h2 | GTAGCCAATTTGGAGAGTGTTCGAGATACCTTGTGGCAGATT<br>AACACACCACAAGGTATCTCGAACACTCTC | | |
| B17 | aaaaa **CGATTGTTTGTTGTGGACGCATGCTAATCGGATGAG** | B17h1 | GTGGACACCTGCTAATCGGATGAGTGTTCGTTATCGCTCATC<br>CGATTAGCAGGTGTCCACAACAAACAATCG | Alexa Fluor 405 | PMID: 38966983 |
| | | B17h2 | CGATAACGAACACTCATCCGATTAGCAGGTGTCCACCGATT<br>GTTTGTTGTGGACACCTGCTAATCGGATGAG | | |

#### Supplemental Table S3: mRNA split initiator oligonucleotide probe sequences used in this study (designed with Özpolat Lab HCR probe generator; PMID: 34793615)

| Target mRNA | Accession no/reference | Split initiator design | Probe name | mRNA probe (italicized) + linker sequence (lowercase) + split initiator sequence (bold) (5'-3') |
|---|---|---|---|---|
| GFP (from p006-GFP-pBAD plasmid) | PMID: 37841098 | B1 | B1-pbadGFP-split-Ia | **GAGGAGGGCAGCAAACGG** aa *AGCAGCTGTTACAAACTCAAGAAGG* |
| | | | B1-pbadGFP-split-Ib | *TAGTTCATCCATGCCATGTGTAATC* ta **GAAGAGTCTTCCTTTACG** |
| | | | B1-pbadGFP-split-IIa | **GAGGAGGGCAGCAAACGG** aa *CATCGCCAATTGGAGTATTTTGTTG* |
| | | | B1-pbadGFP-split-IIb | *AATGGTTGTCTGGTAAAAGGACAGG* ta **GAAGAGTCTTCCTTTACG** |
| | | | B1-pbadGFP-split-IIIa | **GAGGAGGGCAGCAAACGG** aa *GCCATGATGTATACATTGTGTGAGT* |
| | | | B1-pbadGFP-split-IIIb | *GCTTTGATTCCATTCTTTTGTTTGT* ta **GAAGAGTCTTCCTTTACG** |
| | | | B1-pbadGFP-split-IVa | **GAGGAGGGCAGCAAACGG** aa *TGTTTCCATCTTCTTTAAAATCAAC* |
| | | | B1-pbadGFP-split-IVb | *AGTTGTATTCCAATTTGTGTCCAAG* ta **GAAGAGTCTTCCTTTACG** |
| | | | B1-pbadGFP-split-Va | **GAGGAGGGCAGCAAACGG** aa *ATCACCTTCAAACTTGACTTCAGCA* |
| | | | B1-pbadGFP-split-Vb | *TTTTAACTCGATTCTATTAACAAGG* ta **GAAGAGTCTTCCTTTACG** |
| | | | B1-pbadGFP-split-VIa | **GAGGAGGGCAGCAAACGG** aa *AATATAGTTCTTTCCTGTACATAAC* |
| | | | B1-pbadGFP-split-VIb | *GTCTTGTAGTTCCCGTCATCTTTGA* ta **GAAGAGTCTTCCTTTACG** |
| | | | B1-pbadGFP-split-VIIa | **GAGGAGGGCAGCAAACGG** aa *GCCGTTTCATGTGATCTGGGTATCT* |
| | | | B1-pbadGFP-split-VIIb | *CGGGCATGGCACTCTTGAAAAAGTC* ta **GAAGAGTCTTCCTTTACG** |
| | | | B1-pbadGFP-split-VIIIa | **GAGGAGGGCAGCAAACGG** aa *AGTGACAAGTGTTGGCCACGGAACT* |
| | | | B1-pbadGFP-split-VIIIb | *AAAGCATTGAACACCATAAGAGAGA* ta **GAAGAGTCTTCCTTTACG** |
| | | | B1-pbadGFP-split-IXa | **GAGGAGGGCAGCAAACGG** aa *AGGGTAAGTTTTCCGTATGTTGCAT* |
| | | | B1-pbadGFP-split-IXb | *AGTTTTCCAGTAGTGCAAATAAATT* ta **GAAGAGTCTTCCTTTACG** |
| | | | B1-pbadGFP-split-Xa | **GAGGAGGGCAGCAAACGG** aa *TGTGCCCATTAACATCACCGTCTAA* |
| | | | B1-pbadGFP-split-Xb | *CTTCTCCCTCTCCACTGACAGAAAA* ta **GAAGAGTCTTCCTTTACG** |
| mCherry (from SL1344 glmS::mCherry) | PMID: 25121752 | B4 | B4-mCherry split-1a | **CCTCAACCTACCTCCAAC** aa *GATCGCCATATTGTCTTCTTCACCT* |
| | | | B4-mCherry split-1b | *AACTTTGAAACGCATGAACTCTTTG* at **TCTCACCATATTCGCTTC** |
| | | | B4-mCherry split-2a | **CCTCAACCTACCTCCAAC** aa *GGACGACCCTCACCTTCGCCTTCGA* |
| | | | B4-mCherry split-2b | *AGTTTCGCGGTCTGGGTACCTTCGT* at **TCTCACCATATTCGCTTC** |
| | | | B4-mCherry split-3a | **CCTCAACCTACCTCCAAC** aa *TGGAGACAGGATGTCCCACGCGAAC* |
| | | | B4-mCherry split-3b | *GTACGCCTTGCTACCGTACATAAAC* at **TCTCACCATATTCGCTTC** |
| | | | B4-mCherry split-4a | **CCTCAACCTACCTCCAAC** aa *CCGGGAAAGACAGTTTCAGGTAATC* |
| | | | B4-mCherry split-4b | *TCATAACACGTTCCCACTTGAAACC* at **TCTCACCATATTCGCTTC** |
| | | | B4-mCherry split-5a | **CCTCAACCTACCTCCAAC** aa *GAGAAGAGTCCTGGGTAACGGTAAC* |
| | | | B4-mCherry split-5b | *CTTTGTAGATAAATTCACCGTCCTG* at **TCTCACCATATTCGCTTC** |
| | | | B4-mCherry split-6a | **CCTCAACCTACCTCCAAC** aa *TGCATAACCGGACCGTCGCTAGGGA* |
| | | | B4-mCherry split-6b | *GACGCTTCCCAGCCCATCGTTTTTT* at **TCTCACCATATTCGCTTC** |
| | | | B4-mCherry split-7a | **CCTCAACCTACCTCCAAC** aa *ACGCTGTTTGATCTCGCCTTTCAGC* |
| | | | B4-mCherry split-7b | *GTAGTGACCGCCGTCTTTGAGCTTC* at **TCTCACCATATTCGCTTC** |
| | | | B4-mCherry split-8a | **CCTCAACCTACCTCCAAC** aa *CGGGAGCTGAACCGGCTTTTTAGCT* |
| | | | B4-mCherry split-8b | *CAGCTTGATATTAACGTTGTACGCA* at **TCTCACCATATTCGCTTC** |
| | | | B4-mCherry split-9a | **CCTCAACCTACCTCCAAC** aa *CAGCTTGATATTAACGTTGTACGCA* |
| | | | B4-mCherry split-9b | *ATACCACCGGTAGAGTGACGGCCCT* at **TCTCACCATATTCGCTTC** |
| *Stm* OmpA (strain SL1344) | FQ312003 | B1 | B1_OmpA_Ia | **GAGGAGGGCAGCAAACGG** aa *ACGATCCGGAGCCAGGCAATCGAT* |
| | | | B1_OmpA_Ib | *AACGCCTTTAACTTCGATCTCTACG* ta **GAAGAGTCTTCCTTTACG** |
| | | | B1_OmpA_IIa | **GAGGAGGGCAGCAAACGG** aa *GGTGTTGCCGGTAACCGGGTTAGAT* |
| | | | B1_OmpA_IIb | *GGCAGCGCGAGGTTTCACGTTGTCA* ta **GAAGAGTCTTCCTTTACG** |
| | | | B1_OmpA_IIIa | **GAGGAGGGCAGCAAACGG** aa *CAGGTAATCAACAACAGACTGAGCA* |
| | | | B1_OmpA_IIIb | *TTTGTCAGACGGAATACCTTTGGAG* ta **GAAGAGTCTTCCTTTACG** |
| | | | B1_OmpA_IVa | **GAGGAGGGCAGCAAACGG** aa *GTCAGAACCGATACGGTCAGTGAAG* |
| | | | B1_OmpA_IVb | *TTTCTCGGACAGACCCTGGTTGTAA* ta **GAAGAGTCTTCCTTTACG** |
| | | | B1_OmpA_Va | **GAGGAGGGCAGCAAACGG** aa *ATCCAGGTTGCTCAGCTGGCTGTAC* |
| | | | B1_OmpA_Vb | *AGAACGACAACGGAACCGTCTTTC* ta **GAAGAGTCTTCCTTTACG** |
| | | | B1_OmpA_VIa | **GAGGAGGGCAGCAAACGG** aa *TTCAGGGTAGATTTGTTGAAGTTG* |
| | | | B1_OmpA_VIb | *TGATCCAGAGCCTGCTGGCCTTCCG* ta **GAAGAGTCTTCCTTTACG** |
| | | | B1_OmpA_VIIa | **GAGGAGGGCAGCAAACGG** aa *TTGGTCTGTACTTCCGGAGCCGGA* |
| | | | B1_OmpA_VIIb | *AGTACGTCAGACTTCAGAGTGAAGT* ta **GAAGAGTCTTCCTTTACG** |
| | | | B1_OmpA_VIIIa | **GAGGAGGGCAGCAAACGG** aa *TCTTGCTGGCCGAAACGGTAGGAAA* |
| | | | B1_OmpA_VIIIb | *GGTGCCGGAGCTACTACCGGAGCAG* ta **GAAGAGTCTTCCTTTACG** |
| | | | B1_OmpA_IXa | **GAGGAGGGCAGCAAACGG** aa *CGGGTGCCGATGGTGTTGGCATCAC* |
| | | | B1_OmpA_IXb | *CTACGCTCAGCAGGCCGTTGTCCG* ta **GAAGAGTCTTCCTTTACG** |
| | | | B1_OmpA_Xa | **GAGGAGGGCAGCAAACGG** aa *GTAGACGGGCCGCCAGGGACGTTAG* |
| | | | B1_OmpA_Xb | *GGGAAACGCCGGTGTCGTGGTCTT* ta **GAAGAGTCTTCCTTTACG** |
| | | | B1_OmpA_XIa | **GAGGAGGGCAGCAAACGG** aa *CCAGACGGGTATAAACGTCCAGAT* |
| | | | B1_OmpA_XIb | *TGGTGTCTGCACGCCATACCATACC* ta **GAAGAGTCTTCCTTTACG** |
| | | | B1_OmpA_XIIa | **GAGGAGGGCAGCAAACGG** aa *TCAACTGAACGCCCTGAGCTTTTAT* |
| | | | B1_OmpA_XIIb | *CAGTGATTGGATAACCCAGTTTAGC* ta **GAAGAGTCTTCCTTTACG** |
| | | | B1_OmpA_XIIIa | **GAGGAGGGCAGCAAACGG** aa *GCATACGGCCTAACCAGTCGTAGCC* |
| | | | B1_OmpA_XIIIb | *CGCCATTGATGTTGTCGCCTTTGTA* ta **GAAGAGTCTTCCTTTACG** |
| | | | B1_OmpA_XIVa | **GAGGAGGGCAGCAAACGG** aa *ATGAAGCCGGTGTCATGGTACTGAG* |
| | | | B1_OmpA_XIVb | *TTTCATGAGTCGGGCCATCATTGT* ta **GAAGAGTCTTCCTTTACG** |
| | | | B1_OmpA_XVa | **GAGGAGGGCAGCAAACGG** aa *TGTTATCTTTCGGAGCGGCCTGCG* |
| | | | B1_OmpA_XVb | *AGCCCAGTTTAGCACCAGCGTACCA* ta **GAAGAGTCTTCCTTTACG** |

#### Supplemental Table S4: Prospective longitudinal preterm infant cohort study characteristics

| Characteristics | NEC | Control |
|---|---|---|
| Number of patients | 7 | 6 |
| Average gestational age at birth (weeks) | 27 4/7 | 28 1/7 |
| Mean age at NEC diagnosis (DOL) | 41 | N/A |
| Biological sex | Females, *n* = 5<br>Males, *n* = 2 | Females, *n* = 4<br>Males, *n* = 2 |
| Mean birth weight (g) | 1009.7 | 1141.8 |
| Mode of delivery | Vaginal, *n* = 2<br>Caesarean, *n* = 5 | Vaginal, *n* = 2<br>Caesarean, *n* = 4 |
| Feeding modality | MOM/partial MOM, *n* = 5<br>Exclusive DM, *n* = 2 | MOM/partial MOM, *n* = 5<br>Exclusive DM, *n* = 1 |

This prospective longitudinal cohort of preterm infant patients included pre-NEC samples and matched controls (gestational age 24 to 30 weeks) collected prospectively (from DOL 10 to 53) and assigned to groups after NEC diagnosis or discharge from the neonatal intensive care unit.

#### Supplemental Table S5: Bacterial culture conditions and strains

| Bacteria isolate | Source | Growth media | Time of growth |
|---|---|---|---|
| **Facultative anaerobes** | | | |
| *Salmonella enterica serovar Typhimurium* (SL1344) | M. Raffatellu | LB broth | 18 hours |
| *Salmonella enterica serovar Typhimurium* (SL1344-mCherry) | M. Raffatellu<br>L. Knodler | LB broth | 18 hours |
| *Escherichia* coli (DH5α; pBAD-GFP) | Addgene #108315 | LB broth | 18 hours |
| *Lactobacillus gasseri* | Hand, PITT | MRS broth | 48 hours |
| *Streptococcus agalactiae* (BAA-2675) | As previously described (PMID: 37462916) | | |
| *Staphylococcus aureus* (CT1) | | | |
| *Pseudomonas aeruginosa* (01) | | | |
| *Escherichia coli* (605) | | | |
| *Klebsiella oxytoca* (K405) | | | |
| *Klebsiella pneumoniae* | | | |
| *Enterococcus faecalis* (19433) | | | |
| **Obligate anaerobes** | | | |
| *Akkermansia muciniphila* (Muc) | ATCC BAA-835 | Reinforced clostridial medium | 48 hours |
| *Fusobacterium nucleatum subsp. nucleatum* (VPI 4351) | ATCC 23726 | Reinforced clostridial medium | 48 hours |
| *Bifidobacterium breve* (S1) | ATCC 15700 | Bifidus selective medium broth | 48-72 hours |
| *Bifidobacterium longum subsp. infantis* (S12) | ATCC 15697 | Bifidus selective medium broth | 48-72 hours |
| *Bacteroides ovatus* (NCTC 11153) | ATCC 8483 | Peptone yeast extract broth | 24 hours |
| *Clostridium perfringens* | Hand, PITT | Reinforced clostridial medium or peptone yeast extract broth | 24 hours |
| *Clostridioides difficile* (90556-M6S) | ATCC 9689 | Reinforced clostridial medium | 24 hours |
| *Flavonifractor plautii* (VPI 0310) | ATCC 29863 | Reinforced clostridial medium | 24-48 hours |
| *Faecalibacterium prausnitzii* (VPI C13-51) | ATCC 27768 | Reinforced clostridial medium | 72-96 hours |
| *Ruminococcus gnavus* (VPI C7-9) | ATCC 29149 | Reinforced clostridial medium | 48 hours |
| *Finegoldia magna* (WAL2508) | ATCC 29328 | Reinforced clostridial medium | 24 hours |
| *Veillonella parvula* (ATCC 17742) | ATCC 10790 | Reinforced clostridial medium | 48 hours |
| PedsCom isolates:<br>*Anaerostipes sp.*<br>*Clostridium intestinale*<br>*Enterococcus faecalis*<br>*Ligilactobacillus murinus*<br>*Lactobacillus johnsonii*<br>*Kosakonia cowanii*<br>*Parabacteroides distasonis*<br>*Staphylococcus xylosus*<br>*Mammalicoccus sciuri* | As previously described (PMID: 36996818) | | |
